# Biological structure and function emerge from scaling unsupervised learning to 250 million protein sequences

**DOI:** 10.1101/622803

**Authors:** Alexander Rives, Joshua Meier, Tom Sercu, Siddharth Goyal, Zeming Lin, Jason Liu, Demi Guo, Myle Ott, C. Lawrence Zitnick, Jerry Ma, Rob Fergus

**Affiliations:** Facebook AI Research; Dept. of Computer Science, New York University; Harvard University; Booth School of Business, University of Chicago & Yale Law School

## Abstract

In the field of artificial intelligence, a combination of scale in data and model capacity enabled by unsupervised learning has led to major advances in representation learning and statistical generation. In the life sciences, the anticipated growth of sequencing promises unprecedented data on natural sequence diversity. Protein language modeling at the scale of evolution is a logical step toward predictive and generative artificial intelligence for biology. To this end we use unsupervised learning to train a deep contextual language model on 86 billion amino acids across 250 million protein sequences spanning evolutionary diversity. The resulting model contains information about biological properties in its representations. The representations are learned from sequence data alone. The learned representation space has a multi-scale organization reflecting structure from the level of biochemical properties of amino acids to remote homology of proteins. Information about secondary and tertiary structure is encoded in the representations and can be identified by linear projections. Representation learning produces features that generalize across a range of applications, enabling state-of-the-art supervised prediction of mutational effect and secondary structure, and improving state-of-the-art features for long-range contact prediction.

## 1. Introduction

Growth in the number of protein sequences in public databases has followed an exponential trend over decades, creating a deep view into the breadth and diversity of protein sequences across life. This data is a promising ground for studying predictive and generative models for biology using artificial intelligence. Our focus here will be to fit a single model to many diverse sequences from across evolution. Accordingly we study high-capacity neural networks, investigating what can be learned about the biology of proteins from modeling evolutionary data at scale.

The idea that biological function and structure are recorded in the statistics of protein sequences selected through evolution has a long history (Yanofsky et al., 1964; Altschuh et al., 1987; 1988). Out of the possible random perturbations to a sequence, evolution is biased toward selecting those that are consistent with fitness (Göbel et al., 1994). The unobserved variables that determine a protein’s fitness, such as structure, function, and stability, leave a record in the distribution of observed natural sequences (Göbel et al., 1994).

Unlocking the information encoded in protein sequence variation is a longstanding problem in biology. An analogous problem in the field of artificial intelligence is natural language understanding, where the distributional hypothesis posits that a word’s semantics can be derived from the contexts in which it appears (Harris, 1954).

Recently, techniques based on self-supervision, a form of unsupervised learning in which context within the text is used to predict missing words, have been shown to materialize representations of word meaning that can generalize across natural language tasks (Collobert & Weston, 2008; Dai & Le, 2015; Peters et al., 2018; Devlin et al., 2018). The ability to learn such representations improves significantly with larger training datasets (Baevski et al., 2019; Radford et al., 2019).

Protein sequences result from a process greatly dissimilar to natural language. It is uncertain whether the models and objective functions effective for natural language transfer across differences between the domains. We explore this question by training high-capacity Transformer language models on evolutionary data. We investigate the resulting unsupervised representations for the presence of biological organizing principles, and information about intrinsic biological properties. We find metric structure in the representation space that accords with organizing principles at scales from physicochemical to remote homology. We also find that secondary and tertiary protein structure can be identified in representations. The structural properties captured by the representations generalize across folds. We apply the representations to a range of prediction tasks and find that they improve state-of-art features across the applications.

## 2. Background

Sequence alignment and search is a longstanding basis for comparative and statistical analysis of biological sequence data. (Altschul et al., 1990; Altschul & Koonin, 1998; Eddy, 1998; Remmert et al., 2011). Search across large databases containing evolutionary diversity assembles related sequences into a multiple sequence alignment (MSA). Within sequence families, mutational patterns convey information about functional sites, stability, tertiary contacts, binding, and other properties (Altschuh et al., 1987; 1988; Göbel et al., 1994). Conserved sites correlate with functional and structural importance (Altschuh et al., 1987). Local biochemical and structural contexts are reflected in preferences for distinct classes of amino acids (Levitt, 1978). Covarying mutations have been associated with function, tertiary contacts, and binding (Göbel et al., 1994).

The prospect of inferring biological structure and function from evolutionary statistics has motivated development of machine learning on individual sequence families. Direct coupling analysis (Lapedes et al., 1999; Thomas et al., 2008; Weigt et al., 2009) infers constraints on the structure of a protein by fitting a generative model in the form of a Markov Random Field (MRF) to the sequences in the protein’s MSA. Various methods have been developed to fit the MRF (Morcos et al., 2011; Jones et al., 2011; Balakrishnan et al., 2011; Ekeberg et al., 2013b). The approach can also be used to infer functional constraints (Figliuzzi et al., 2016; Hopf et al., 2017), and the generative picture can be extended to include latent variables (Riesselman et al., 2018).

Recently, self-supervision has emerged as a core direction in artificial intelligence research. Unlike supervised learning which requires manual annotation of each datapoint, self-supervised methods use unlabeled datasets and thus can exploit far larger amounts of data. Self-supervised learning uses proxy tasks for training, such as predicting the next word in a sentence given all previous words (Bengio et al., 2003; Dai & Le, 2015; Peters et al., 2018; Radford et al., 2018; 2019) or predicting words that have been masked from their context (Devlin et al., 2018; Mikolov et al., 2013).

Increasing the dataset size and the model capacity has shown improvements in the learned representations. In recent work, self-supervision methods used in conjunction with large data and high-capacity models produced new state-of-the-art results approaching human performance on various question answering and semantic reasoning benchmarks (Devlin et al., 2018), and coherent natural text generation (Radford et al., 2019).

This paper explores self-supervised language modeling approaches that have demonstrated state-of-the-art performance on a range of natural language processing tasks, applying them to protein data in the form of unlabeled amino acid sequences. Since protein sequences use a small vocabulary of twenty canonical elements, the modeling problem is more similar to character-level language models (Mikolov et al., 2012; Kim et al., 2016) than word-level models. Like natural language, protein sequences also contain long-range dependencies, motivating use of architectures that detect and model distant context (Vaswani et al., 2017).

## 3. Scaling language models to 250 million diverse protein sequences

Large protein sequence databases contain diverse sequences sampled across life. In our experiments we explore datasets with up to 250 million sequences of the Uniparc database (The UniProt Consortium, 2007) which has 86 billion amino acids. This data is comparable in size to large text datasets that are being used to train high-capacity neural network architectures on natural language (Devlin et al., 2018; Radford et al., 2019). To model the data of evolution with fidelity, neural network architectures must have capacity and inductive biases to represent its breadth and diversity.

We investigate the Transformer (Vaswani et al., 2017), which has emerged as a powerful general-purpose model architecture for representation learning and generative modeling, outperforming recurrent and convolutional architectures in natural language settings. We use a deep Transformer (Devlin et al., 2018), taking as input amino acid character sequences.

The Transformer processes inputs through a series of blocks that alternate self-attention with feed-forward connections. Self-attention allows the network to build up complex representations that incorporate context from across the sequence. Since self-attention explicitly constructs pairwise interactions between all positions in the sequence, the Transformer architecture directly represents residue-residue interactions.

We train models using the masked language modeling objective (Devlin et al., 2018). Each input sequence is corrupted by replacing a fraction of the amino acids with a special mask token. The network is trained to predict the missing tokens from the corrupted sequence:

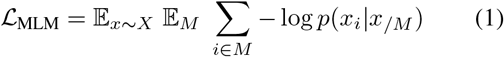

For each sequence *x* we sample a set of indices *M* to mask, replacing the true token at each index *i* with the mask token. For each masked token, we independently minimize the negative log likelihood of the true amino acid *x*_*i*_ given the masked sequence *x*_*/M*_ as context. Intuitively, to make a prediction for a masked position, the model must identify dependencies between the masked site and the unmasked parts of the sequence.

### Evaluation of language models

We begin by training a series of Transformers on all the sequences in UniParc (The UniProt Consortium, 2007), holding out a random sample of 1M sequences for validation. We use these models throughout to investigate properties of the representations and the information learned during pre-training.

To comparatively evaluate generalization performance of different language models we use UniRef50 (Suzek et al., 2015), a clustering of UniParc at 50% sequence identity. For evaluation, a held-out set of 10% of the UniRef50 clusters is randomly sampled. The evaluation dataset consists of the representative sequences of these clusters. All sequences belonging to the held-out clusters are removed from the pre-training datasets.

We explore the effect of the underlying sequence diversity in the pre-training data. Clustering UniParc shows a power-law distribution of cluster sizes (Suzek et al., 2007), implying the majority of sequences belong to a small fraction of clusters. Training using a clustering of the sequences results in a re-weighting of the masked language modeling loss toward a more diverse set of sequences. We use UniRef (Suzek et al., 2015) to create three pre-training datasets with differing levels of diversity: (i) the low-diversity dataset (UR100) uses the UniRef100 representative sequences; (ii) the high-diversity sparse dataset (UR50/S) uses the UniRef50 representative sequences; (iii) the high-diversity dense dataset (UR50/D) samples the UniRef100 sequences evenly across the UniRef50 clusters.

Table 1 presents modeling performance on the held-out UniRef50 sequences across a series of experiments exploring different model classes, number of parameters, and pre-training datasets. Models are compared using the exponentiated cross entropy (ECE) metric, which is the exponential of the model’s loss averaged per token. In the case of the Transformer this is 2 ^*ℒ* MLM^. ECE describes the mean uncertainty of the model among its set of options for every prediction: ranging from 1 for an ideal model to 25 (the number of unique amino acid tokens in the data) for a completely random prediction. To measure the difficulty of generalization to the evaluation set, we train a series of n-gram models across a range of context lengths and settings of Laplace smoothing on UR50/S. The best n-gram model has an ECE of 17.18 with context size of 4.

**Table 1.** Evaluation of language models for generalization to held-out UniRef50 clusters. (a) Exponentiated cross-entropy (ECE) ranges from 25 for a random model to 1 for a perfect model. (b) Best n-gram model across range of context sizes and Laplace-smoothing settings. (c) State-of-the-art LSTM bidirectional language models (Peters et al., 2018). (d) Transformer model baselines with 6 and 12 layers. Small Transformer models have better performance than LSTMs despite having fewer parameters. (e) 34-layer Transformer models trained on datasets of differing sequence diversity. Increasing the diversity of the training set improves generalization. High-capacity Transformer models outperform LSTMs and smaller Transformers. (f) 34-layer Transformer models trained on reduced fractions of data. Increasing training data improves generalization.

|  | **Model** |  | **Params** | **Training** | **ECE** |
| --- | --- | --- | --- | --- | --- |
| (a) | Oracle |  |  |  | 1 |
|  | Uniform Random |  |  |  | 25 |
| (b) | n-gram | 4-gram |  | UR50/S | 17.18 |
| (c) | LSTM | Small | 28.4M | UR50/S | 14.42 |
|  | LSTM | Large | 113.4M | UR50/S | 13.54 |
| (d) | Transformer | 6-layer | 42.6M | UR50/S | 11.79 |
|  | Transformer | 12-layer | 85.1M | UR50/S | 10.45 |
| (e) | Transformer | 34-layer | 669.2M | UR100 | 10.32 |
|  | Transformer | 34-layer | 669.2M | UR50/S | 8.54 |
|  | Transformer | 34-layer | 669.2M | UR50/D | **8.46** |
| (f) | Transformer | 10% data | 669.2M | UR50/S | 10.99 |
|  | Transformer | 1% data | 669.2M | UR50/S | 15.01 |
|  | Transformer | 0.1% data | 669.2M | UR50/S | 17.50 |

As a baseline we train recurrent LSTM bidirectional language models (Peters et al., 2018), which are state-of-the-art for recurrent models in the text domain. Unlike standard left-to-right autoregressive LSTMs, these models use the whole sequence context, making them comparable to the Transformers we study. We evaluate a small model with approximately 25M parameters, and a large model with approximately 113M parameters. Trained on the UR50/S dataset, the small and large LSTM models have an ECE of 14.4 and 13.5 respectively.

We also train two small Transformers, a 12-layer (85.1M parameters) and 6-layer Transformer (42.6M parameters) on the UR50/S dataset. Both Transformer models have better ECE values (10.45, and 11.79 respectively) than the small and large LSTM models, despite the large LSTM having more parameters. These results show the Transformer enables higher fidelity modeling of protein sequences for a comparable number of parameters.

We train high-capacity 34-layer Transformers (approx 670M parameters) across the three datasets of differing diversity. The high-capacity Transformer model trained on the UR50/S dataset outperforms the smaller Transformers indicating an improvement in language modeling with increasing model capacity. Transformers trained on the two high-diversity datasets, UR50/S and UR50/D, improve generalization over the UR100 low-diversity dataset. The best Transformer trained on the most diverse and dense dataset reaches an ECE of 8.46, meaning that intuitively the is model choosing among approximately 8.46 amino acids for each prediction.

We also train a series of 34-layer Transformer models on 0.1%, 1%, and 10% of the UR50/S dataset, seeing the expected relationship between increased data and improved generalization performance. Underfitting is observed even for the largest models trained on 100% of UR50/S suggesting potential for additional improvements with higher capacity models.

### ESM-1b Transformer

Finally we perform a systematic optimization of model hyperparameters on 100M parameter models to identify a robust set of hyperparameters. The hyperparameter search is described in detail in Appendix B. We scale the hyperparameters identified by this search to train a model with approximately 650M parameters (33 layers) on the UR50/S dataset, resulting in the ESM-1b Transformer.

## 4. Multi-scale organization in sequence representations

The variation observed in large protein sequence datasets is influenced by processes at many scales, including properties that affect fitness directly, such as activity, stability, structure, binding, and other properties under selection (Hormoz, 2013; Hopf et al., 2017) as well as by contributions from phylogenetic bias (Gabaldon, 2007), experimental and selection biases (Wang et al., 2019; Overbaugh & Bangham, 2001), and sources of noise such as random genetic drift (Kondrashov et al., 2003).

Unsupervised learning may encode underlying factors that, while unobserved, are useful for explaining the variation in sequences seen by the model during pre-training. We investigate the representation space of the network at multiple scales from biochemical to evolutionary homology to look for signatures of biological organization.

Neural networks contain inductive biases that impart structure to representations. Randomly initialized networks can produce features that perform well without any learning (Jarrett et al., 2009). To understand how the process of learning shapes the representations, it is necessary to compare representations before and after they have been trained. Furthermore, a basic level of intrinsic organization is expected in the sequence data itself as a result of biases in amino acid composition. To disentangle the role of frequency bias in the data we also compare against a baseline that maps each sequence to a vector of normalized amino acid counts.

### Learning encodes biochemical properties

The Transformer neural network represents the identity of each amino acid in its input and output embeddings. The input embeddings project the input amino acid tokens into the first Transformer block. The output embeddings project the final hidden representations back to logarithmic probabilities. The interchangeability of amino acids within a given structural or functional context in a protein depends on their biochemical properties (Hormoz, 2013). Self-supervision can be expected to capture these patterns to build a representation space that reflects biochemical knowledge.

To investigate if the network has learned to encode physicochemical properties in representations, we project the weight matrix of the final embedding layer of the network into two dimensions with t-SNE (Maaten & Hinton, 2008). In Figure 1 the structure of the embedding space reflects biochemical interchangeability with distinct clustering of hydrophobic and polar residues, aromatic amino acids, and organization by molecular weight and charge.

**Figure 1.**
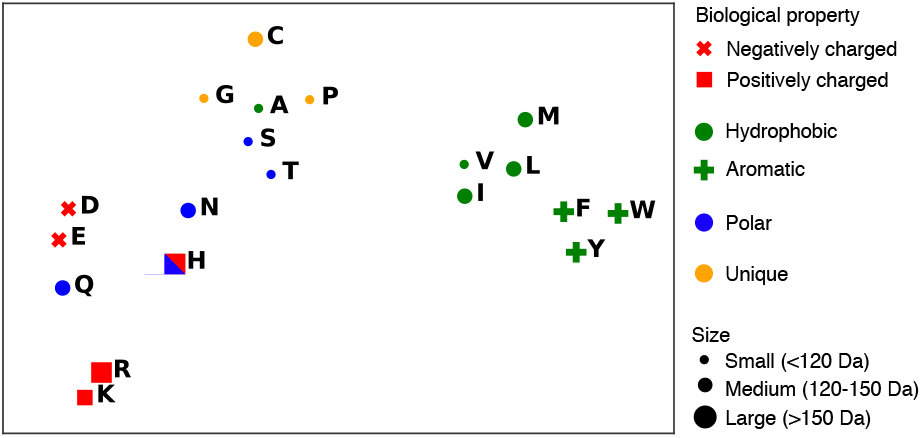
Biochemical properties of amino acids are represented in the Transformer model’s output embeddings, visualized here with t-SNE. Through unsupervised learning, residues are clustered into hydrophobic, polar, and aromatic groups, and reflect overall organization by molecular weight and charge. Visualization of 36-layer Transformer trained on UniParc.

### Biological variations are encoded in representation space

Each protein can be represented as a single vector by averaging across the hidden representation at each position in its sequence. Protein embeddings represent sequences as points in a high dimensional space. Each sequence is represented as a single point and sequences assigned to similar representations by the network are mapped to nearby points. We investigate how homologous genes are represented in this space.

The structure and function of orthologous genes is likely to be retained despite divergence of their sequences (Huerta-Cepas et al., 2018). We find in Figure 2a that training shapes the representation space so that orthologous genes are clustered. Figure 2a shows a two-dimensional projection of the model’s representation space using t-SNE. Prior to training the organization of orthologous proteins in the model’s representation space is diffuse. Orthologous genes are clustered in the learned representation space.

**Figure 2.**
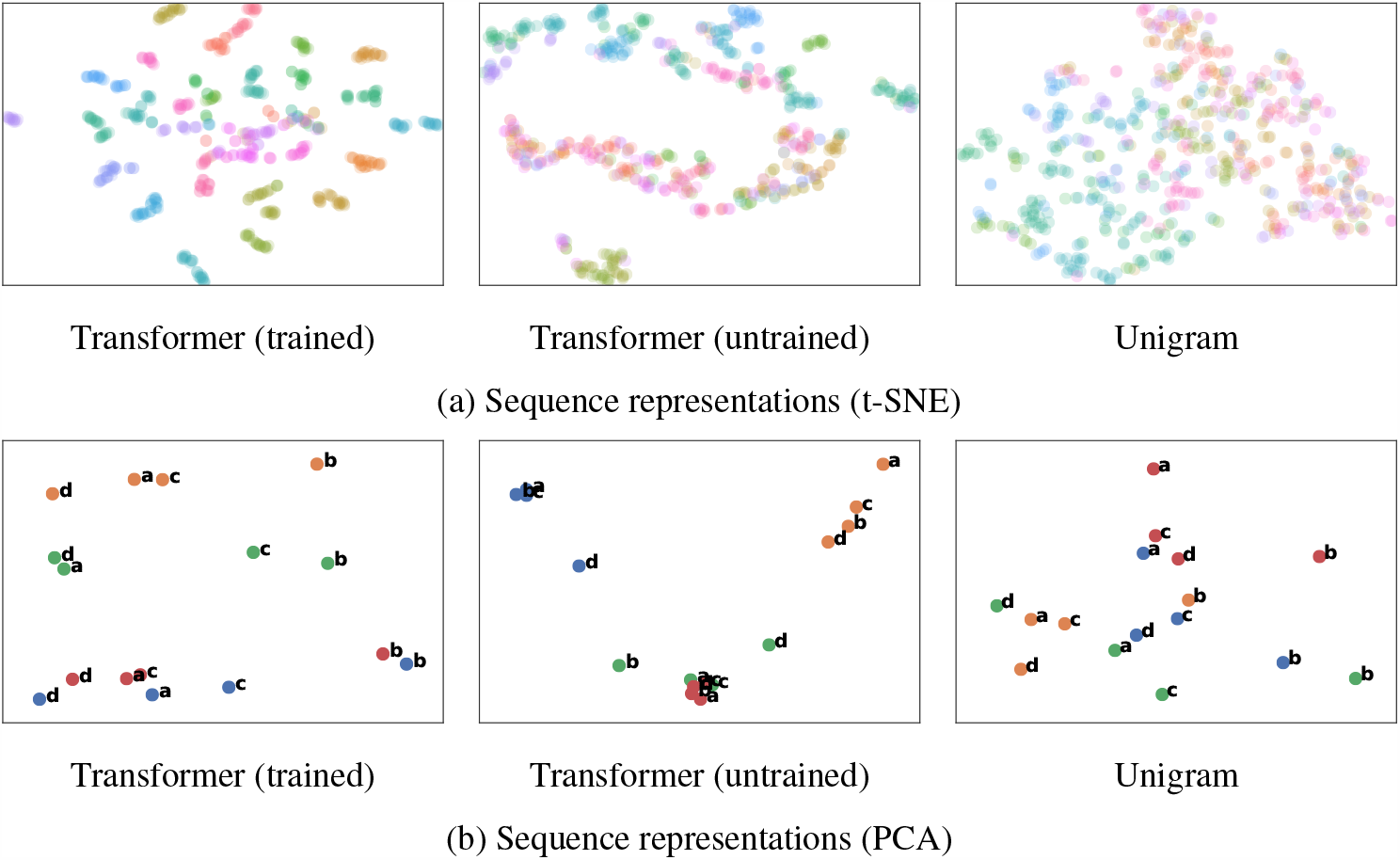
Protein sequence representations encode and organize biological variations. (a) Each point represents a gene, and each gene is colored by the orthologous group it belongs to (dimensionality is reduced by t-SNE). Orthologous groups of genes are densely clustered in the trained representation space. By contrast, the untrained representation space and unigram representations do not reflect strong organization by evolutionary relationships. (b) Genes corresponding to a common biological variation are related linearly in the trained representation space. Genes are colored by their orthologous group, and their species is indicated by a character label. PCA recovers a species axis (horizontal) and orthology axis (vertical) in the trained representation space, but not in the untrained or unigram spaces. Representations are from the 36-layer Transformer model trained on UniParc.

We examine whether unsupervised learning encodes biological variations into the structure of the representation space. We apply principal component analysis (PCA), to recover principal directions of variation in the representations, selecting 4 orthologous genes across 4 species to look for directions of variation. Figure 2b indicates that linear dimensionality reduction recovers species and orthology as primary axes of variation in the representation space after training. This form of structure is absent from the representations prior to training.

To quantitatively investigate the structure of the representation space, we assess nearest neighbor recovery under vector similarity queries. If biological properties are encoded along independent directions in the representation space, then proteins corresponding with a unique biological variation are related by linear vector arithmetic. In Figure S1 we find that learning improves recovery of target proteins under queries encoded as linear transformations along the species or gene axes.

### Learning encodes remote homology

Remotely homologous proteins have underlying structural similarity despite divergence of their sequences. If structural homology is encoded in the metric structure of the representation space, then the distance between proteins reflects their degree of structural relatedness.

We investigate whether the representation space enables detection of remote homology at the superfamily (proteins that belong to different families but are in the same superfamily) and fold (proteins that belong to different superfamilies but have the same fold) level. We construct a dataset to evaluate remote homology detection using SCOPe (Fox et al., 2014). Following standard practices (Söding & Remmert, 2011) we exclude Rossman-like folds (c.2-c.5, c.27 and 28, c.30 and 31) and four-to eight-bladed *β*-propellers (b.66-b.70).

An unsupervised classifier on distance from the query measures the density of homologous proteins in the neighborhood of a query sequence. For each domain, a vector similarity query is performed against all other domains, ranking them by distance to the query domain. For evaluation at the fold level, any domain with the same fold is a positive; any domain with a different fold is a negative; and domains belonging to the same superfamily are excluded. For evaluation at the superfamily level, any domain with the same superfamily is a positive; any domain with a different superfamily is a negative; and domains belonging to the same family are excluded. We report AUC for the classifier, and Hit-10 (Ma et al., 2014) which gives the probability of recovering a remote homolog in the ten highest ranked results.

Table 2 indicates that vector nearest neighbor queries using the representations can detect remote homologs that are distant at the fold level with similar performance to HH-blits (Remmert et al., 2011) a state-of-the-art HMM-HMM alignment-based method. At the superfamily level, where sequence similarity is higher, HMM performance is better, but Transformer embeddings are close. Fast vector nearest neighbor finding methods allow billions of sequences to be searched for similarity to a query protein within milliseconds (Johnson et al., 2017).

**Table 2.**
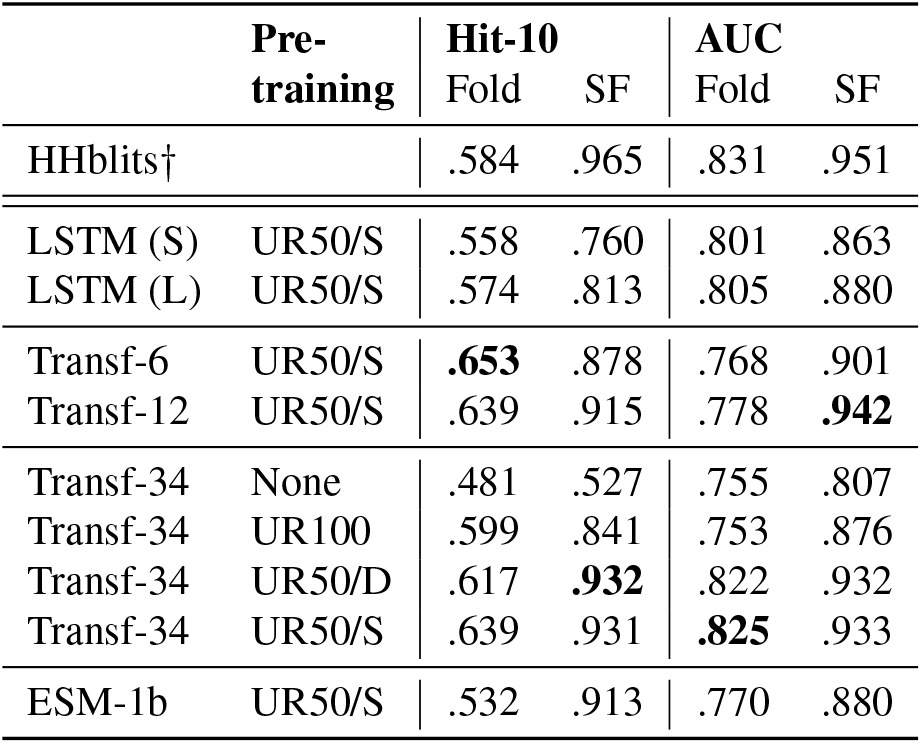
Remote homology detection. Structural homology at the fold and superfamily (SF) level is encoded in the metric structure of the representation space. Results for unsupervised classifier based on distance between vector sequence embeddings. Hit-10 reports the probability that a remote homolog is included in the ten nearest neighbors of the query sequence. Area under the ROC curve (AUC) is reported for classification by distance from the query in representation space. Transformer models have higher performance than LSTMs and similar performance to HMMs at the fold level. Best neural models are indicated in bold. *†*HH-blits (Remmert et al., 2011), a state-of-the-art HMM-based method for remote homology detection, using 3 iterations of sequence search.

### Learning encodes alignment within a protein family

An MSA identifies corresponding sites across a family of related sequences (Ekeberg et al., 2013a). These correspondences give a picture of evolutionary variation at different sites within the sequence family. The model receives as input individual sequences and is given no access to the family of related sequences except via learning. We investigate whether the final hidden representations of a sequence encode information about the family it belongs to.

Family information could appear in the network through assignment of similar representations to positions in different sequences that are aligned in the family’s MSA. Using the collection of MSAs of structurally related sequences in Pfam (Bateman et al., 2013), we compare the distribution of cosine similarities of representations between pairs of residues that are aligned in the family’s MSA to a background distribution of cosine similarities between unaligned pairs of residues. A large difference between the aligned and unaligned distributions implies that the representations use shared features for related sites within all the sequences of the family.

Figure 3a depicts the distribution of cosine similarity values between aligned and unaligned positions within a representative family for the trained model and baselines. Unsupervised learning produces a marked shift between the distributions of aligned and unaligned pairs. Figure 3b and Figure 3c indicate that these trends hold under the constraints that the residue pairs (1) share the same amino acid identity or (2) have different amino acid identities.

**Figure 3.**
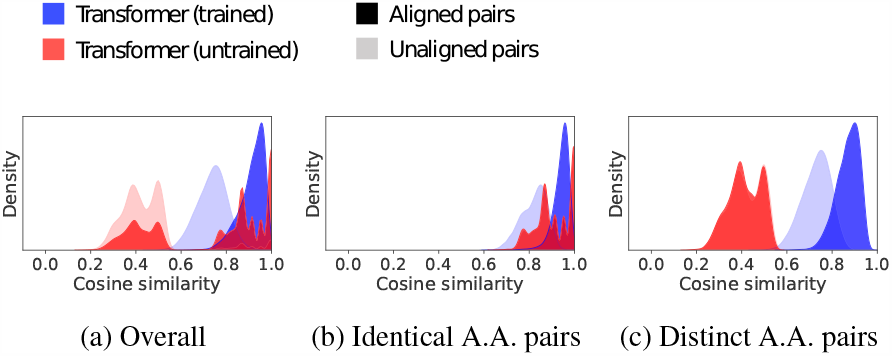
Final representations from trained models implicitly align sequences. Cosine similarity distributions are depicted for the final representations of residues from sequences within PFAM family PF01010. The differences between the aligned (dark blue) and unaligned (light blue) distributions imply that the trained Transformer representations are a powerful discriminator between aligned and unaligned positions in the sequences. In contrast representations prior to training do not separate the aligned (dark red) and unaligned positions (light red). AUCs across 128 PFAM families are reported in Table S1.

We estimate differences between the aligned and unaligned distributions across 128 Pfam families using the area under the ROC curve (AUC) as a metric of discriminative power between aligned and unaligned pairs. Table S1 shows a quantitative improvement in average AUC after unsupervised training, supporting the idea that self-supervision encodes information about the MSA of a sequence into its representation of the sequence.

## 5. Prediction of secondary structure and tertiary contacts

There is reason to believe that unsupervised learning will cause the model’s representations to contain structural information. The underlying structure of a protein is a hidden variable that influences the patterns observed in sequence data. For example local sequence variation depends on secondary structure (Levitt, 1978); and tertiary structure introduces higher order dependencies in the choices of amino acids at different sites within a protein (Marks et al., 2011; Anishchenko et al., 2017). While the model cannot observe protein structure directly, it observes patterns in the sequences of its training data that are determined by structure. In principle, the network could compress sequence variations by capturing commonality in structural elements across the data, thereby encoding structural information into the representations.

### Linear projections

We begin by identifying information about protein structure that is linearly encoded within the representations. The use of linear projections ensures that the information originates in the Transformer representations, enabling a direct inspection of the structural content of representations. By comparing representations of the Transformer before and after pre-training, we can identify the information that emerges as a result of the unsupervised learning.

We perform a five-fold cross validation experiment to study generalization of structural information at the family, superfamily, and fold level. For each of the three levels, we construct a dataset of 15,297 protein structures using the SCOPe database. We partition the structures into five parts, splitting by family, superfamily, and fold accordingly. Fivefold cross validation is performed independently for each of the levels of structural hold-out.

To detect information about secondary structure we fit a logistic regression to the hidden representations using the 8-class secondary structure labels. To detect information about tertiary structure, we fit two separate linear projections to the hidden representations of pairs of positions in the sequence, taking their dot product to regress a binary variable indicating whether the positions are in contact in the protein’s 3-dimensional structure. The neural representations are compared to (i) projections of the sequence profile; (ii) unsupervised contacts predicted by the CCMpred implementation (Seemayer et al., 2014) of direct coupling analysis. MSAs for the baselines are generated from the UniClust30 (Mirdita et al., 2017) database using 3 iterations of search by Hhblits. For secondary structure, we report 8-class accuracies. For contact precision we report Top-L long-range precision, i.e. the precision of the L (length of the protein) highest ranked predictions for contacts with sequence separation of at least 24 residues.

Table 3 shows results of the cross validation experiments. Prior to pre-training, minimal information about secondary structure and contacts can be detected. After pre-training, projections recover information about secondary structure and long-range contacts that generalizes across families, superfamilies, and folds. Secondary structure prediction 8-class accuracy distributions (Figure S2), and long-range contact prediction Top-L precision distributions (Figure S3) demonstrate that pre-training produces an increase in structural information across the entire distribution of test domains. Table 3 shows that projections of the Transformer representations recover more structure than projections of the sequence profile. For long-range contacts, projections of the best Transformer models have higher precision than contacts predicted by CCMpred across all levels of structural generalization. As the level of structural split becomes more remote, there is little degradation for secondary structure, with performance at the family level similar to the fold level. For long-range contacts, while generalization is reduced at the fold level in comparison to the family level, the best models still capture more structure than the unsupervised baseline. Training with higher-diversity sequences (UR50 datasets) improves learning of both secondary structure and long-range contacts, with a more pronounced effect on longrange contacts.

**Table 3.** Linear projections. Five-fold cross validation experiment for generalization at the family, superfamily, and fold level. 8-class accuracy (secondary structure), Top-L long-range precision (contacts), mean and standard deviation across test sets for the five partitions. Minimal information about structure is present in representations prior to training. Information about secondary and tertiary structure emerges in representations as a result of unsupervised learning on sequences with the language modeling objective. Increasing diversity of sequences improves learning of structure. (Higher diversity UR50 datasets improve over UR100). Learned representations enable linear projections to generalize to held-out folds, outperforming projections of the sequence profile, and contacts identified by the CCMpred (Seemayer et al., 2014) implementation of direct coupling analysis.

| Model | Pre-training | SSP |  |  | Contact |  |  |
| --- | --- | --- | --- | --- | --- | --- | --- |
|  |  | Family | Superfamily | Fold | Family | Superfamily | Fold |
| HMM Profile | - | $47.9 \pm 1.1$ | $47.8 \pm 1.3$ | $48.0 \pm 1.6$ | $14.8 \pm 0.6$ | $14.5 \pm 1.4$ | $14.6 \pm 1.6$ |
| CCMpred | - | - | - | - | $32.7 \pm 1.0$ | $32.5 \pm 2.3$ | $32.6 \pm 0.4$ |
| Transformer-6 | UR50/S | $62.4 \pm 0.7$ | $61.8 \pm 0.7$ | $61.8 \pm 1.0$ | $24.4 \pm 2.2$ | $19.4 \pm 3.1$ | $18.6 \pm 0.5$ |
| Transformer-12 | UR50/S | $65.5 \pm 0.7$ | $65.0 \pm 1.0$ | $65.2 \pm 1.0$ | $32.8 \pm 2.9$ | $25.8 \pm 3.7$ | $25.2 \pm 0.9$ |
| Transformer-34 | (None) | $41.5 \pm 0.9$ | $41.3 \pm 0.8$ | $41.3 \pm 1.3$ | $8.7 \pm 0.3$ | $8.4 \pm 0.7$ | $8.4 \pm 0.3$ |
| Transformer-34 | UR100 | $65.4 \pm 0.7$ | $64.9 \pm 0.9$ | $64.9 \pm 0.9$ | $28.3 \pm 2.5$ | $22.5 \pm 3.3$ | $22.0 \pm 0.8$ |
| Transformer-34 | UR50/S | $69.4 \pm 0.6$ | $68.8 \pm 0.9$ | $68.9 \pm 0.9$ | $43.9 \pm 2.8$ | $36.4 \pm 4.2$ | $35.3 \pm 1.7$ |
| Transformer-34 | UR50/D | $69.5 \pm 0.6$ | $68.9 \pm 0.9$ | $69.0 \pm 0.9$ | $43.9 \pm 2.8$ | $37.1 \pm 4.6$ | $36.3 \pm 2.0$ |
| ESM-1b | UR50/S | **$71.1 \pm 0.5$** | **$70.5 \pm 0.9$** | **$70.6 \pm 0.9$** | **$49.2 \pm 2.5$** | **$43.5 \pm 4.8$** | **$42.8 \pm 2.3$** |

Figure 4 visualizes three-class secondary structure projections for two domains belonging to held-out folds. Prior to pre-training, projections produce an incoherent prediction of secondary structure. After pre-training projections recover a coherent prediction with most errors occurring at the boundaries of secondary structure regions. Figure 5 compares a projected contact map to predictions from CCMpred. Transformer projections recover complex contact patterns, including long-range contacts. Further visualizations of projected contacts for eight randomly selected test proteins are shown in Figure S7.

**Figure 4.**
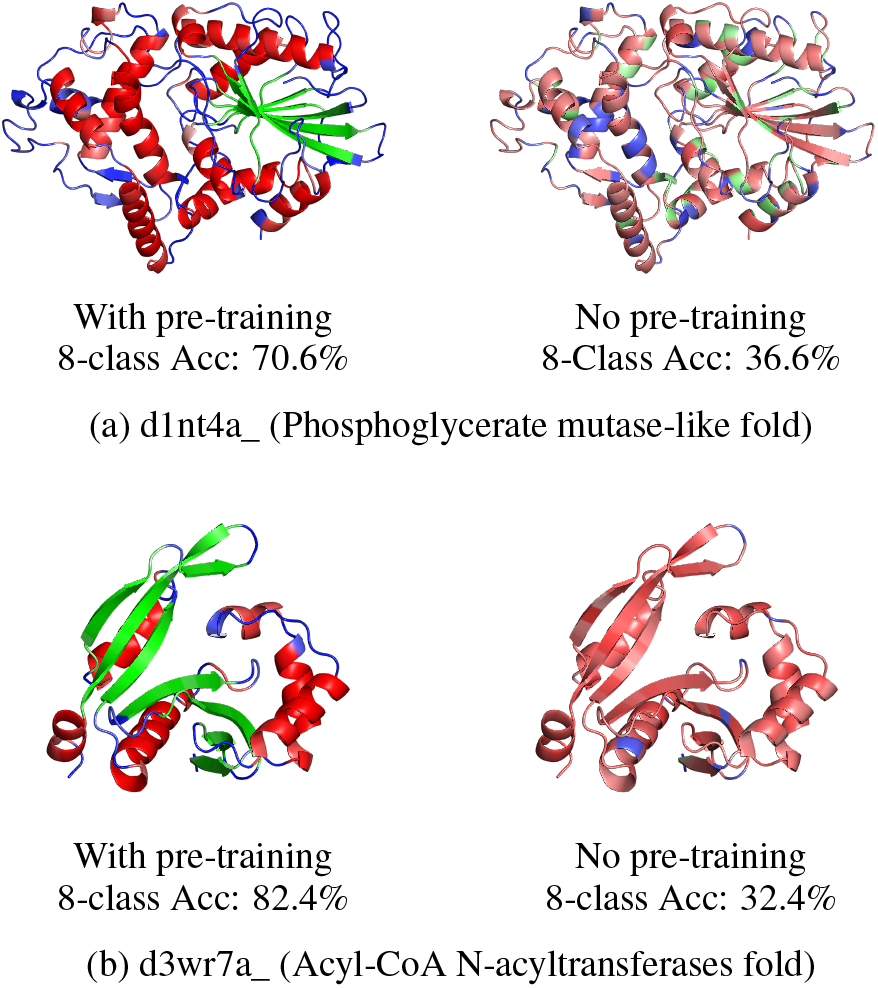
Secondary structure (linear projections). Example predictions for held-out folds. Unsupervised pre-training encodes secondary structure into representations. Following pre-training, linear projections recover secondary structure (left column). With-out pre-training little information is recovered (right column). Colors indicate secondary structure class identified by the projection: helix (red), strand (green), and coil (blue). Color intensities indicate confidence. Representations from ESM-1b Transformer are used.

**Figure 5.**
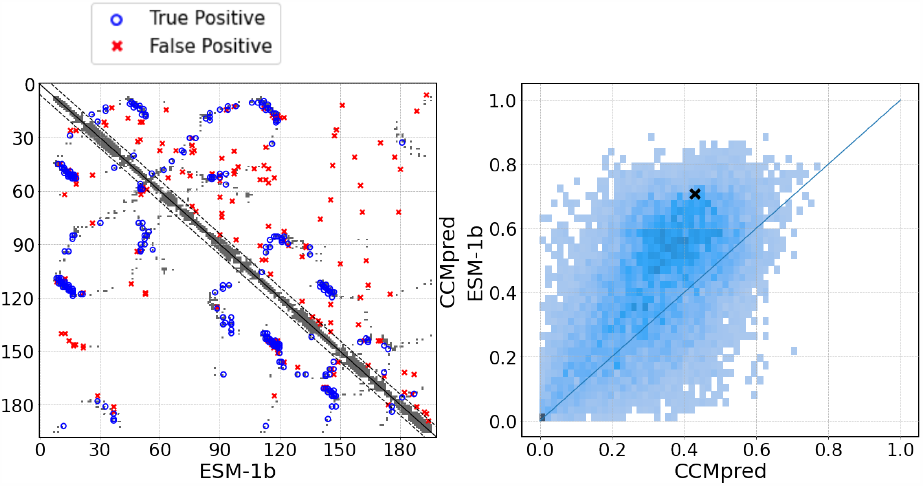
Residue-residue contacts (linear projections). Left: Top-L predictions for fold level held-out example d1n3ya_, with vWA-like fold. True positives in blue, false positives in red, superimposed on ground-truth contact map in grey. ESM-1b Transformer projections below the diagonal, CCMpred predictions above the diagonal. Right: Precision distribution (Top-L long-range) comparing ESM-1b projections with CCMpred across all domains in the five test partitions with structural hold-out at the fold level. Visualized domain marked by x.

### Deep neural network

We train deep neural networks to predict secondary structure and contacts from the representations. We choose state-of-the-art neural architectures for both tasks. These downstream models are trained with a supervised loss to predict either the secondary structure or contact map from the pre-trained representations. The architecture of the downstream model is kept fixed across experiments with different representations and baselines to enable comparison.

To predict secondary structure we replace the linear layer with a deep neural network, using the model architecture introduced by the Netsurf method (Klausen et al., 2019). For tertiary structure, we predict the binary contact map from the hidden representation of the sequence. We use a dilated convolutional residual network similar to recent state-of-the-art methods for tertiary structure prediction (Xu, 2018; Jones & Kandathil, 2018; Senior et al., 2018).

Table 4 compares the representations for secondary structure prediction. We evaluate models on the CB513 test set (Cuff & Barton, 1999) and the CASP13 domains (Kryshtafovych et al., 2019). For comparison we also re-implement the Netsurf method. The models are trained on the Netsurf training dataset which applies a 25% sequence identity holdout with CB513, and a temporal hold-out with CASP13. The Transformer features are compared before and after unsupervised pre-training to features from the LSTM baselines. They are also compared to the HMM profiles used by Netsurf. The best Transformer features (71.6%) match the performance of the HMM profiles (71.2%), and exceed the published performance of RaptorX (70.6)% on the same benchmark (Klausen et al., 2019), implying that protein langauge models can produce features that are directly competitive with sequence profiles for secondary structure prediction.

**Table 4.**
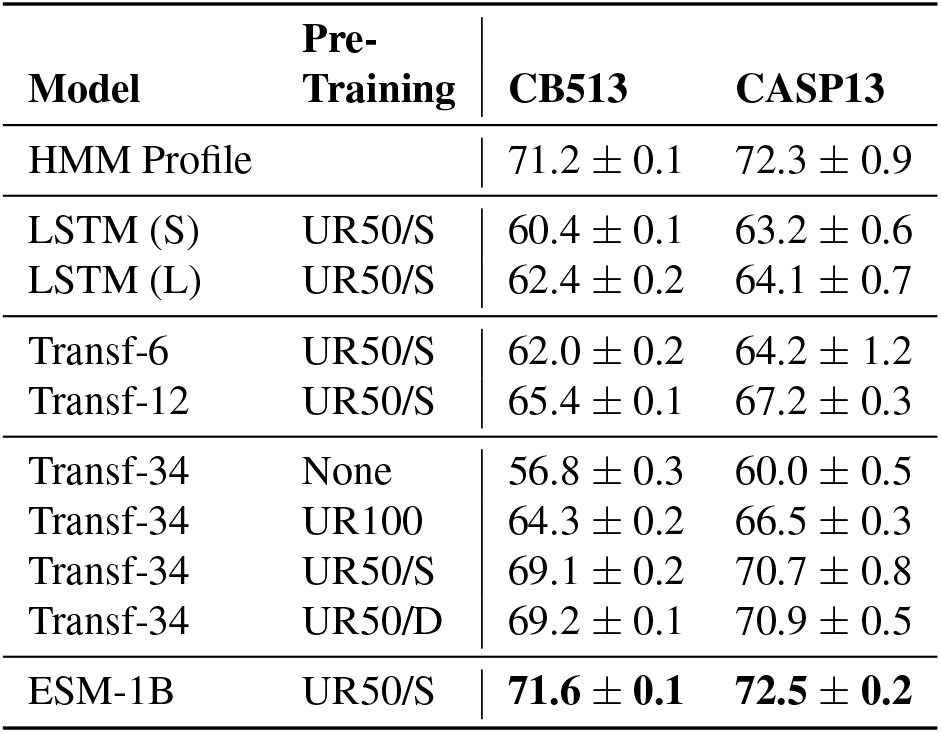
Eight-class secondary structure prediction accuracy on the CB513 and CASP13 test sets. A fixed neural architecture is trained to predict the secondary structure label from the language model representation of the input sequence. The Transformer has higher performance than the comparable LSTM baselines. Pretraining with the high-diversity UR50 datasets increases accuracy significantly. Features from ESM-1b Transformer are competitive with HMM profiles for supervised secondary structure prediction.

| Model | Pre-Training | CB513 | CASP13 |
| --- | --- | --- | --- |
| HMM Profile | | 71.2 ± 0.1 | 72.3 ± 0.9 |
| LSTM (S) | UR50/S | 60.4 ± 0.1 | 63.2 ± 0.6 |
| LSTM (L) | UR50/S | 62.4 ± 0.2 | 64.1 ± 0.7 |
| Transf-6 | UR50/S | 62.0 ± 0.2 | 64.2 ± 1.2 |
| Transf-12 | UR50/S | 65.4 ± 0.1 | 67.2 ± 0.3 |
| Transf-34 | None | 56.8 ± 0.3 | 60.0 ± 0.5 |
| Transf-34 | UR100 | 64.3 ± 0.2 | 66.5 ± 0.3 |
| Transf-34 | UR50/S | 69.1 ± 0.2 | 70.7 ± 0.8 |
| Transf-34 | UR50/D | 69.2 ± 0.1 | 70.9 ± 0.5 |
| ESM-1B | UR50/S | **71.6 ± 0.1** | **72.5 ± 0.2** |

Table 5 shows performance of the various representations for predicting Top-L long-range contacts across a panel of benchmarks using the RaptorX train set (Wang et al., 2017). For comparison we train the same architecture using features from RaptorX (Wang et al., 2017; Xu, 2018). The Test (Wang et al., 2017) and CASP11 (Moult et al., 2016) test sets evaluate with sequence identity hold-out at 25%; the CASP12 (Moult et al., 2018) test set implements a temporal hold-out with the structural training data; and the CASP13 (Kryshtafovych et al., 2019) experiment implements a full temporal hold-out of both the pre-training and training data. For contact prediction, the best features from representation learning do not achieve comparable performance to the state-of-the-art RaptorX features (50.2 vs 59.4 respectively on the RaptorX test set).

**Table 5.** Top-L long-range contact precision. A deep dilated convolutional residual network is trained to predict contacts using the representations from the pre-trained language model. The pre-trained Transformer representations outperform the LSTM representations in all cases. Pre-training on the high-diversity UR50 datasets boosts precision of representations over pre-training on UR100. High-capacity Transformers (34 layer) outperform lower capacity models (6/12 layer).

| Model | Pre-training | Test | CASP |  |  |
| --- | --- | --- | --- | --- | --- |
|  |  |  | 11 | 12 | 13 |
| LSTM (S) | UR50/S | 24.1 | 23.6 | 19.9 | 15.3 |
| LSTM (L) | UR50/S | 27.8 | 26.4 | 24.0 | 16.4 |
| Transf-6 | UR50/S | 30.2 | 29.9 | 25.3 | 19.8 |
| Transf-12 | UR50/S | 37.7 | 33.6 | 27.8 | 20.7 |
| Transf-34 | (None) | 16.3 | 17.7 | 14.8 | 13.3 |
| Transf-34 | UR100 | 32.7 | 28.9 | 24.3 | 19.1 |
| Transf-34 | UR50/S | 50.2 | 42.8 | 34.7 | 30.1 |
| Transf-34 | UR50/D | 50.0 | 43.0 | 33.6 | 28.0 |
| ESM-1b | UR50/S | **56.9** | **47.4** | **42.7** | **35.9** |

In the secondary structure benchmarks Transformer representations produce higher accuracy than the LSTM baselines with comparable numbers of parameters. For contact prediction Transformer reprentations yield higher precision than LSTMs, with even the smallest Transformer representations exceeding LSTMs with more parameters. Diversity in the pre-training data also has a strong effect, with the high-diversity datasets providing significant improvements over the low-diversity dataset. Relative performance of the representations is consistent across all four of the contact benchmarks using different hold-out methodology.

### Relationship between language modeling and structure learning

To investigate the relationship between the language modeling objective and information about structure in the model, linear projections for secondary structure and contacts are fit using the representations from Transformer models taken from checkpoints across their pre-training trajectories. We use the Transformers trained on UR50/S. We fit the projections and evaluate with the train and test split implemented by the first partition of the fold level structural hold-out dataset. For each model, Figure 6 shows a linear relationship between the language modeling objective and information about structure, which is maintained over the course of pre-training. The linear fit is close to ideal for both secondary structure and contacts. A similar experiment is also performed for secondary structure with a deep neural network instead of linear projection, using the Netsurf training sequences and CB513 test set. A linear relationship between secondary structure accuracy and language modeling ECE is also observed for the deep prediction head (Figure S4). Thus, for a given model and pre-training dataset, language modeling fidelity measured by ECE is a good proxy for the structural content of the representations. Since performance on the language modeling objective improves with model capacity, this suggests further scale may improve results on structure prediction tasks.

**Figure 6.**
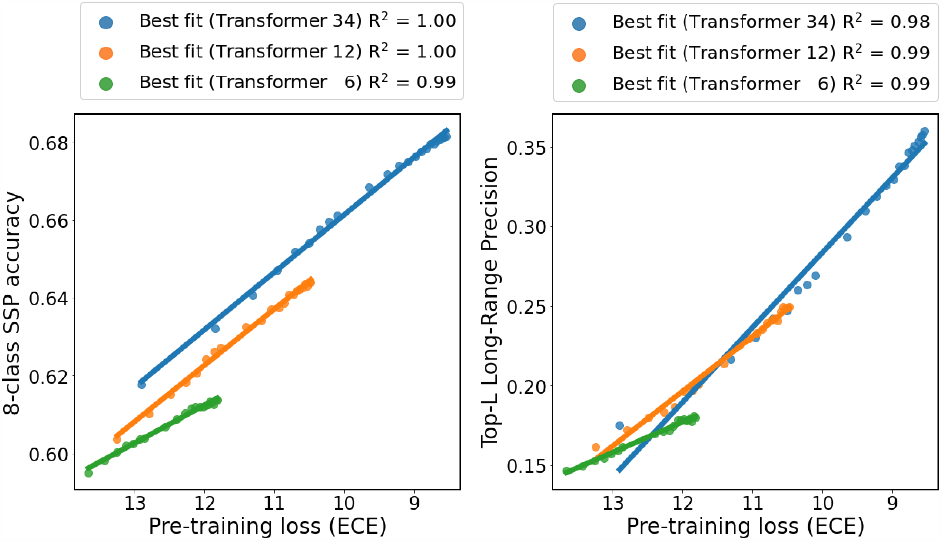
Relationship between the language modeling objectiveand structure learning. Eight-class secondary structure predictionaccuracy (left) and contact prediction Top-L long-range precision(right) both as a function of pre-training ECE. Performance isevaluated on held-out folds. Linear projections are fit using modelcheckpoints over the course of pre-training on UR50/S. The linearrelationship for each model indicates that for a given model and pre-training dataset, the language modeling ECE is a good proxy for the structural content of the representations. Improvement ofthe model’s ECE leads to an increase in information about structure.This establishes a link between the language modeling objectiveand unsupervised structure learning.

### Single versus multi-family pre-training

We compare training across evolutionary statistics to training on single protein families. We pre-train separate 12-layer Transformer models on the Pfam multiple sequence alignments of the three most common domains in nature longer than 100 amino acids, the ATP-binding domain of the ABC transporters, the protein kinase domain, and the response regulator receiver domain. We test the ability of models trained on one protein family to generalize secondary structure information within-family and out-of-family by evaluating on sequences with ground truth labels from the family the model was trained on or from the alternate families. The models are evaluated using linear projections. In all cases, the model trained on within-family sequences has higher accuracy than models trained on out-of-family sequences (Table S2), indicating poor generalization when training on single MSA families. More significantly, the model trained across the full UniParc sequence diversity has a higher accuracy than the single-family model accuracies, even on the same-family evaluation dataset. This suggests that the representations learned from the full dataset are generalizing information about secondary structure learned outside the sequence family.

## 6. Feature combination

Features discovered by unsupervised protein language modeling can be combined with state-of-the-art features to improve them further. Current state-of-the-art methods use information derived from MSAs. We combine this information with features from the Transformer model.

We explore three approaches for incorporating information from representation learning. For each input sequence *x*: (i) *direct* uses the final hidden representation from the Transformer directly; (ii) *avg* takes the average of the final hidden representation at each position across the sequences from the MSA of *x*; (iii) *cov* produces features for each pair of positions, using the uncentered covariance across sequences from the MSA of *x*, after dimensionality reduction of the final hidden representations by PCA. Note that (i) and (ii) produce features for each position in *x*, while (iii) produces features for each pair of positions.

### Secondary structure

Current state-of-the-art methods for secondary structure prediction have high accuracies for the eight-class prediction problem. We investigate whether performance can be improved by combining Transformer features with sequence profiles. Table 6 shows that combining the representations with profiles further boosts accuracy, resulting in state-of-the-art performance on secondary structure prediction.

**Table 6.** Feature combination (secondary structure prediction).Eight-class accuracy. The language model improves state-of-theartfeatures for secondary structure prediction. Features from areimplementation of Netsurf (Klausen et al., 2019) are combinedwith 34-layer Transformer (UR50/S) embeddings using a two layerBiLSTM architecture. (a) Performance of Netsurf features alone.(b) Direct adds the Transformer representation of the input sequence.(c) Avg adds the average of Transformer features for eachposition in the MSA of the input sequence. Results exceed thosefor state-of-the-art methods RaptorX (70.6%) and Netsurf (72.1%)on the CB513 test set, and for Netsurf (74.0%) on the CASP13evaluation set used here.

| Features | CB513 | CASP13 |
| --- | --- | --- |
| RaptorX | 70.6 |  |
| Netsurf | 72.1 | 74 |
| (a) Netsurf (reimpl.) | 71.2 ± 0.1 | 72.3 ± 0.9 |
| (b) +direct | 72.1 ± 0.1 | 72.2 ± 0.5 |
| (c) +avg | **73.7 ± 0.2** | **75.1 ± 0.4** |

We establish a baseline of performance by reimplementing the Klausen et al. (2019) method using the same features, resulting in an accuracy of 71.2% (vs. published performance of 72.1%) on the CB513 test set. Then we add the the Transformer features using the *direct* and *avg* combination methods; these achieve 0.9% and 2.5% absolute improvement in accuracy respectively. This suggests that the Transformer features contain information not present in the MSA-derived features.

### Residue-residue contacts

Deep neural networks have enabled recent breakthroughs in the prediction of protein contacts and tertiary structure (Xu, 2018; Senior et al., 2018). State-of-the-art neural networks for tertiary structure and contact prediction use deep residual architectures with two-dimensional convolutions over pairwise feature maps to output a contact prediction or distance potential for each pair of residues (Wang et al., 2017; Xu, 2018; Senior et al., 2018).

A variety of input features, training datasets, and supervision signals are used in state-of-the-art methods. To make a controlled comparison, we fix a standard architecture, training dataset, multiple sequence alignments, and set of base input features for all experiments, to which we add pre-trained features from the Transformer model. For the base features we use the RaptorX feature set which includes PSSM, 3-state secondary structure prediction, one-hot embedding of sequence, APC corrected Potts model couplings, mutual information, pairwise contact potential, and predicted accessibility. RaptorX was the winning method for contact prediction in the CASP12 and CASP13 competitions (Xu, 2018). The training and evaluation sets are the same as used in the previous section.

Table 7 indicates that addition of Transformer features from the 34-layer model trained on UR50/S consistently produces an improvement across the test sets. The table shows Top-L long-range precisions reporting mean and standard deviation over 5 different model seeds. *Direct* gives a modest improvement on some test sets. *Avg* improves over direct, and *cov* provides further gains. For example, *cov* produces an absolute improvement of 3.9% on the RaptorX test set, and 1.8% improvement on the CASP13 test set evaluated with temporal hold-outs on both fine-tuning and pre-training data. Additional results and metrics for contact prediction are reported in Table S3.

**Table 7.** Feature combination (contact prediction). Top-L long-range contact precision. The language model improves state-of-the-artfeatures for contact prediction. A deep ResNet with fixed architecture is trained on each feature set to predict binary contacts. (a)performance of state-of-the-art RaptorX (Xu, 2018) features including PSSM, predicted secondary structure, predicted accessibility,pairwise APC-corrected Potts model couplings and mutual information, and a pairwise contact potential. (b) Adds Transformerrepresentation of the input sequence to the feature set. (c) Adds the average Transformer representation at each position of the MSA. (d)Adds the uncentered covariance over the MSA of a low-dimenensional projection of the Transformer features. Features are from the34-layer Transformer pre-trained on UR50/S.

|  | Test | CASP11 | CASP12 | CASP13 |
| --- | --- | --- | --- | --- |
| # domains | 500 | 105 | 55 | 34 |
| (a) RaptorX | 59.4 ± .2 | 53.8 ± .3 | 51.1 ± .2 | 43.4 ± .4 |
| (b) +direct | 61.7 ± .4 | 55.0 ± .1 | 51.5 ± .5 | 43.7 ± .4 |
| (c) +avg | 62.9 ± .4 | 56.6 ± .4 | 52.4 ± .5 | 44.8 ± .8 |
| (d) +cov | **63.3 ± .2** | **56.8 ± .2** | **53.0 ± .3** | **45.2 ± .5** |

## 7. Prediction of mutational effects

The mutational fitness landscape provides deep insight into biology. Coupling next generation sequencing with a mutagenesis screen allows parallel readout of tens of thousands of variants of a single protein (Fowler & Fields, 2014). The detail and coverage of these experiments provides a view into the mutational fitness landscape of individual proteins, giving quantitative relationships between sequence and protein function. We adapt the Transformer protein language model to predict the quantitative effect of mutations.

First we investigate intra-protein variant effect prediction, where a limited sampling of mutations is used to predict the effect of unobserved mutations. This setting has utility in protein engineering applications (Yang et al., 2019). We evaluate the representations on two deep mutational scanning datasets used by recent state-of-the-art methods for variant effect prediction, Envision (Gray et al., 2018) and DeepSequence (Riesselman et al., 2018). Collectively the data includes over 700,000 variant effect measurements from over 100 large-scale experimental mutagenesis datasets.

Fine-tuning the Transformer yields a mutational effect predictor that is comparable to the results of Envision. Envision (Gray et al., 2018) relies on protein structural and evolutionary features to generalize. We assess whether the Transformer can achieve similar generalization results, without direct access to structural features. The same methodology for partitioning data for training and evaluation is used as in Gray et al. (2018) to allow a comparison of the results. We use the 34-layer Transformer trained on UR50/S. Figure 7 shows the fine-tuned Transformer exceeds the performance of Envision on 10 of the 12 proteins. For each protein a fraction *p* = 0.8 of the data is used for training and the remaining data is used for testing. We report mean and standard deviations for five-fold cross-validation in Table S5. Results varying the fraction of data that is used for training are reported in Figure S5.

**Figure 7.**
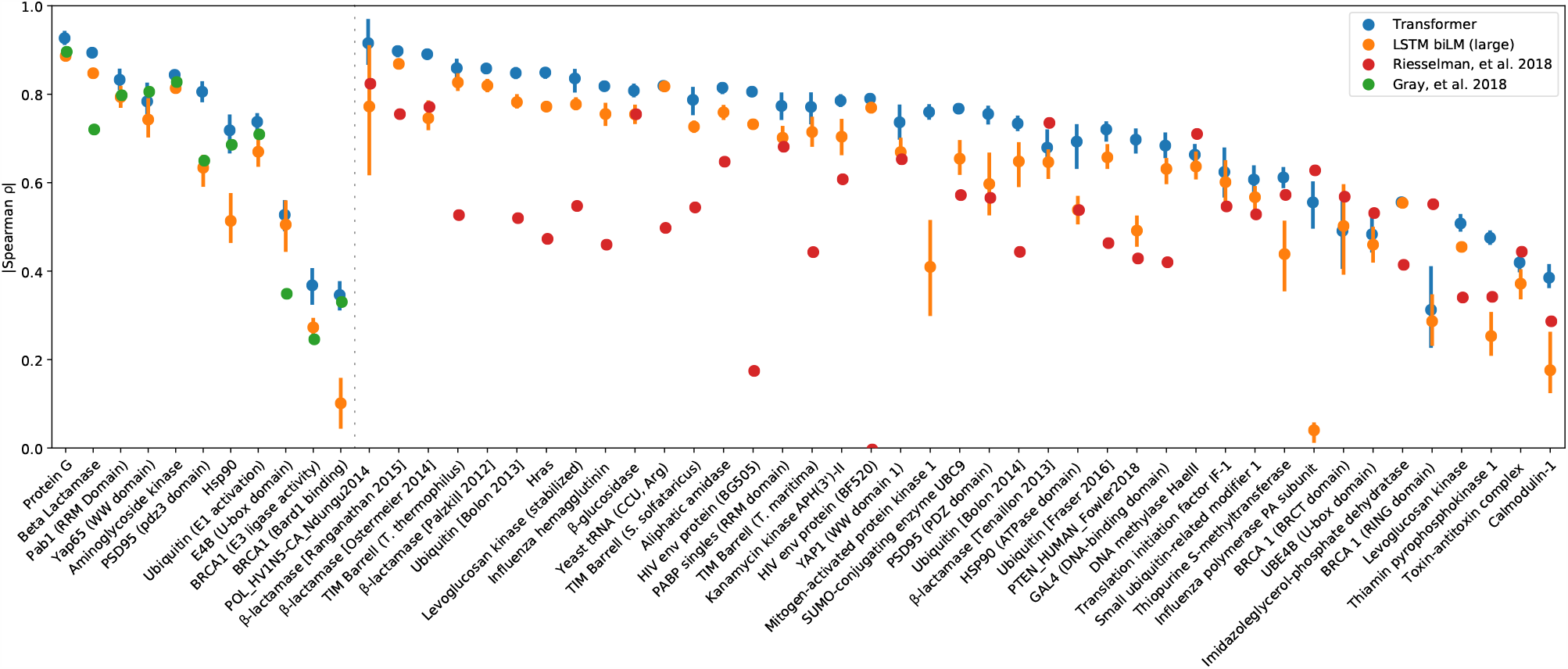
Representation learning enables state-of-the-art supervised prediction of the quantitative effect of mutations. Left panel: Envisiondataset (Gray et al., 2018); right panel: DeepSequence dataset (Riesselman et al., 2018). Transformer representations (34-layer, UR50/S)are compared to the LSTM bidirectional language model (large model, UR50/S). The result of five-fold cross validation is reportedfor each protein. For each partition, supervised fine-tuning is performed on 80% of the mutational data for the protein, and results areevaluated on the remaining 20%. Transformer representations outperform baseline LSTM representations on both datasets. State-of-the-artmethods are also shown for each dataset. Gray et al. (2018) is a supervised method using structural, evolutionary, and biochemicalfeatures, trained with the same protocol as used for the Transformer. Riesselman et al. (2018) is an unsupervised method trained on theMSA of each protein.

We also evaluate using the same five-fold cross validation methodology on the deep mutational scanning experiments assembled for DeepSequence (Riesselman et al., 2018). The fine-tuned Transformer model outperforms the fine-tuned LSTM baselines. While not directly comparable, we also include the performance of the original DeepSequence method which is unsupervised, and represents state-of-the-art for this dataset.

### Generalization to a new fitness landscape

We analyze the Transformer’s ability to generalize to the fitness landscape of a new protein. Following the protocol introduced in Envision, we use a leave-one-out analysis: to evaluate performance on a given protein, we train on data from the remaining *n* − 1 proteins and test on the held-out protein. Figure S6 shows that the Transformer’s predictions from raw sequences perform better than Envision on 5 of the 9 tasks.

## 8. Related Work

Contemporaneously with the preprint of this work, Rives et al. (2019), related preprints Alley et al. (2019) and Heinzinger et al. (2019), also proposed protein language modeling, albeit at a smaller scale. These works, along with Rao et al. (2019), evaluated on a variety of downstream tasks. Rives et al. (2019) first proposed protein language modeling with Transformers. Alley et al. (2019) and Heinzinger et al. (2019) train LSTMs on UniRef50. Rao et al. (2019) trained a 12-layer Transformer model (38M parameters) on Pfam (Bateman et al., 2013). The baselines in this paper are comparable to these models. The large Transformer models trained in this paper are considerably larger than in these related works.

We benchmark against related work in Table 8. Heinzinger et al. (2019), Alley et al. (2019), and Rao et al. (2019), evaluate models on differing downstream tasks and test sets. We retrieve the weights for the above models, evaluating them directly in our codebase against the panel of test sets used in this paper for remote homology, secondary structure prediction, and contact prediction, with the same training data and model architectures. This allows a direct comparison between the representations. Table 8 shows that high-capacity Transformers have strong performance for secondary structure and contact prediction significantly exceeding Alley et al. (2019), Heinzinger et al. (2019), and Rao et al. (2019). The small Transformer models trained as baselines also have higher performance than the methods with comparable parameter numbers.

**Table 8.** Comparison to related protein language models. (RH) Remote Homology at the fold level, using Hit-10 metric on SCOP. (SSP)Secondary structure Q8 accuracy on CB513. (Contact) Top-L long range contact precision on RaptorX test set from Wang et al. (2017).Results for additional test sets in Table S6. ^1^Alley et al. (2019) 2Heinzinger et al. (2019) ^3^Rao et al. (2019). †The pre-training datasets for related work have differences from ours.

| Model | Pre-Training | Params | RH | SSP | Contact |
| --- | --- | --- | --- | --- | --- |
| UniRep<sup>1†</sup> |  | 18M | .527 | 58.4 | 21.9 |
| SeqVec<sup>2†</sup> |  | 93M | .545 | 62.1 | 29.0 |
| Tape<sup>3†</sup> |  | 38M | .581 | 58.0 | 23.2 |
| LSTM (S) | UR50/S | 28.4M | .558 | 60.4 | 24.1 |
| LSTM (L) | UR50/S | 113.4M | .574 | 62.4 | 27.8 |
| Transformer-6 | UR50/S | 42.6M | **.653** | 62.0 | 30.2 |
| Transformer-12 | UR50/S | 85.1M | .639 | 65.4 | 37.7 |
| Transformer-34 | UR100 | 669.2M | .599 | 64.3 | 32.7 |
| Transformer-34 | UR50/S | 669.2M | .639 | 69.2 | 50.2 |
| ESM-1b | UR50/S | 652.4M | .532 | **71.6** | **56.9** |

Protein sequence embeddings have been the subject of recent investigation for protein engineering (Yang et al., 2018). Bepler & Berger (2019) pre-trained LSTMs on protein sequences, adding supervision from contacts to produce embeddings. Subsequent to our preprint, related works have built on its exploration of protein sequence modeling, exploring generative models (Riesselman et al., 2019; Madani et al., 2020), internal representations of Transformers (Vig et al., 2020), and applications of representation learning and generative modeling such as classification (Elnaggar et al., 2019; Strodthoff et al., 2020), mutational effect prediction (Luo et al., 2020), and design of sequences (Repecka et al., 2019; Hawkins-Hooker et al., 2020; Amimeur et al., 2020).

## 9. Discussion

One of the goals for artificial intelligence in biology could be the creation of controllable predictive and generative models that can read and generate biology in its native language. Accordingly, research will be necessary into methods that can learn intrinsic biological properties directly from protein sequences, which can be transferred to prediction and generation.

We investigated deep learning across evolution at the scale of the largest protein sequence databases, training contextual language models across 86 billion amino acids from 250 million sequences. The space of representations learned from sequences by high-capacity networks reflects biological structure at multiple levels, including that of amino acids, proteins, and evolutionary homology. Information about secondary and tertiary structure is internalized and represented within the network. Knowledge of intrinsic biological properties emerges without supervision — no learning signal other than sequences is given during pre-training.

We find that networks that have been trained across evolutionary data generalize: information can be extracted from representations by linear projections, deep neural networks, or by adapting the model using supervision. Fine-tuning produces results that match state-of-the-art on variant activity prediction. Predictions are made directly from the sequence, using features that have been automatically learned by the language model, rather than selected by domain knowledge.

We find that pre-training discovers information that is not present in current state-of-the-art features. The learned features can be combined with features used by state-of-the-art structure prediction methods to improve results. Empirically we find that features discovered by larger models perform better on downstream tasks. The Transformer out-performs LSTMs with similar capacity across benchmarks. Increasing diversity of the training data results in significant improvements to the representations.

While the protein language models we study are of comparable scale to those used in the text domain, our experiments have not yet reached the limit of scale. We observed that even the highest capacity models we trained (with approximately 650-700M parameters) under-fit the sequence datasets, due to insufficient model capacity. The relationship we find between language modeling fidelity and the information about structure encoded into the representations suggests that higher capacity models will yield better representations. These findings imply potential for further model scale and data diversity incorporating sequences from metagenomics.

Combining high-capacity generative models with gene synthesis and high throughput characterization can enable generative biology. The models we have trained can be used to generate new sequences (Wang & Cho, 2019). If neural networks can transfer knowledge learned from protein sequences to design functional proteins, this could be coupled with predictive models to jointly generate and optimize sequences for desired functions. The size of current sequence data and its projected growth point toward the possibility of a general purpose generative model that can condense the totality of sequence statistics, internalizing and integrating fundamental chemical and biological concepts including structure, function, activity, localization, binding, and dynamics, to generate new sequences that have not been seen before in nature but that are biologically active.

### Pre-trained models

Transformer models and baselines are available at: https://github.com/facebookresearch/esm

## Acknowledgments

We thank Tristan Bepler, Richard Bonneau, Yilun Du, Vladimir Gligorijevic, Anika Gupta, Omer Levy, Ian Peikon, Hetunandan Kamisetty, Laurens van der Maaten, Ethan Perez, Oded Regev, Neville Sanjana, and Emily Wang for feedback on the manuscript and insightful conversations. We thank Jinbo Xu for sharing RaptorX features and help with CASP13. We thank Michael Klausen for providing Netsurf training code. Alexander Rives was supported at NYU by NSF Grant #1339362.

## A. Approach & Data

### 1. Pre-training datasets

#### UniParc pre-training dataset

A series of development models are trained on UniParc (The UniProt Consortium, 2007) Release 2019_01 which contains approximately 250M sequences. 1M sequences are held-out randomly for validation. These models were used in the preprint of this paper, and representations from the models are used in Figures 1, 2, and 3.

#### UniRef pre-training datasets

Datasets are based on UniRef (Suzek et al., 2015) dated March 28, 2018 to permit a temporal hold-out with CASP13. 10% of UniRef50 clusters are randomly selected as a held-out evaluation set, yielding 3.02 million representative sequences for evaluation. Three training datasets are used, removing all sequences belonging to clusters selected for the evaluation set: (i) UR100, 124.9M UniRef100 representative sequences; (ii) UR50/S, 27.1M UniRef50 representative sequences; (iii) UR50/D, 124.9M UniRef50 cluster members sampled evenly by cluster. To ensure a deterministic validation set, we removed sequences longer than 1024 amino acids from the validation set.

### 2. Downstream tasks

#### Remote Homology

A dataset of remote homolog pairs is derived from SCOPe (Fox et al., 2014) containing 256,806 pairs of remote homologs at the fold level and 92,944 at the superfamily level, consisting of 217 unique folds and 366 unique superfamilies. Creation of the dataset is detailed below in the section on remote homology.

#### Linear projections

Five-fold cross validation datasets implementing structural hold-outs at the family, superfamily, and fold level are constructed using SCOPe (Fox et al., 2014). Independently for each level of structural hold-out, the domains are split into 5 equal sets, i.e. five sets of folds, superfamilies, or families. This ensures that for each of the five partitions, structures having the same classification do not appear in both the train and test sets. For a given classification level each structure appears in a test set once, so that in the cross validation experiment each of the structures will be evaluated exactly once. Scores reported are the mean and standard deviation over each of the five test sets. Further details on construction of the dataset are given below in the section on linear projections.

#### Secondary structure prediction

All downstream models are trained using the Netsurf (Klausen et al., 2019) training dataset containing 10,837 examples with labels and HMM profiles. Netsurf features are replicated for CASP13 domains using MMseqs2 (Steinegger & Söding, 2017) on the Uniclust90 (Mirdita et al., 2017) dataset released April 2017. For test sets we use (i) the standard CB513 (Cuff & Barton, 1999) test set of 513 sequences with sequence identity hold-out at 25% identity; and (ii) the 34 publicly available CASP domains, using DSSP (Kabsch & Sander, 1983) to label secondary structure, with temporal hold-out for both pre-training and downstream data.

#### Contact prediction

All downstream models are trained using the training and test sets of Wang et al. (2017). Comparisons with RaptorX features use features from Wang et al. (2017) and Xu (2018). The following test sets are used: (i) RaptorX Test, 500 domains (25% sequence identity holdout); (ii) CASP11, 105 domains (25% sequence identity hold-out); (iii) CASP12, 55 domains (temporal hold-out from training data but not pre-training data); (iv) CASP13, 34 publicly released domains (temporal hold-out from training data and pre-training data). The training set consists of 6,767 sequences with contact map targets, a subset of PDB created in February 2015 (Wang et al., 2017). The use of an earlier version of the PDB ensures a temporal holdout w.r.t. both CASP12 and CASP13. Additionally, Wang et al. (2017) implemented a sequence identity hold-out for Test and CASP11 by removing proteins from the training set which share >25% sequence identity or have BLAST E-value <0.1 with the proteins in these test sets.

#### Mutational effect prediction

The model is fine-tuned on deep mutational scanning datasets compiled by Gray et al. (2018) and Riesselman et al. (2018).

### 3. The Transformer

We use a deep Transformer encoder model (Vaswani et al., 2017b; Devlin et al., 2018b), processing input as character sequences of amino acids. In contrast to recurrent and convolutional neural networks, the Transformer makes no assumptions on the ordering of the input and instead uses position embeddings. Particularly relevant to protein sequences is the Transformer’s ability to model long range dependencies, which are not effectively captured by RNNs or LSTMs (Khandelwal et al., 2018). A key factor affecting the performance of LSTMs on these tasks is the path lengths that must be traversed by forward activation and backward gradient signals in the network (Hochreiter et al., 2001).

It is well known that structural properties of protein sequences are reflected in long-range dependencies. Direct coupling analysis (Lapedes et al., 1999; Thomas et al., 2008; Weigt et al., 2009) which aims to detect pairwise dependencies in multiple sequence alignments uses a Markov Random Field (Potts Model) which models the complete sequence with pairwise coupling parameters. Similarly, the Transformer builds up a representation of a sequence by alternating self-attention with non-linear projections. Self-attention structures computation so that each position is represented by a weighted sum of the other positions in the sequence. The attention weights are computed dynamically and allow each position to choose what information from the rest of the sequence to integrate at every computation step.

Developed to model large contexts and long range dependencies in language data, self-attention architectures currently give state-of-the-art performance on various natural language tasks, mostly due to the Transformer’s scalability in parameters and the amount of context it can integrate (Devlin et al., 2018b). The tasks include token-level tasks like part-of-speech tagging, sentence-level tasks such as textual entailment, and paragraph-level tasks like question-answering.

#### Scaled dot-product attention

Self-attention takes a sequence of vectors (*h*_1_, …, *h*_*n*_) and produces a sequence of vectors 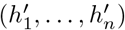 by computing interactions between all elements in the sequence. The Transformer model uses scaled dot-product attention (Vaswani et al., 2017b):

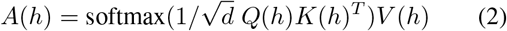

Here the query *Q*, key *K*, and value *V*, are projections of the input sequence to *n* × *d* matrices where *n* is the length of the sequence and *d* is the inner dimension of the matrix outer product between *Q* and *K*. This outer product parameterizes an *n* × *n* map of attention logits, which are rescaled, and passed through the softmax function row-wise, thereby representing each position of the sequence in the output as a convex combination of the sequence of values *V*. One step of self-attention directly models possible pairwise interactions between all positions in the sequence simultaneously. Note the contrast to recurrent and convolutional models which can only represent long-range context through many steps, and the parallel in inductive bias with the explicit pairwise parameterization of Markov Random Fields in widespread use for modeling protein MSAs.

Multi-headed self-attention concatenates the output of *t* independent attention heads:

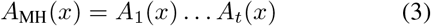

Use of multiple heads enables representation of different inter-position interaction patterns.

#### Architecture

The Transformer models (Vaswani et al., 2017b) in this work take a sequence of tokens (*x*_1_, …, *x*_*n*_) and output a sequence of log probabilities (*y*_1_, …, *y*_*n*_) which are optimized using the masked language modeling objective. The computation proceeds through a series of residual blocks producing hidden states, each a sequence of vectors (*h*_1_, …, *h*_*n*_) with embedding dimension *d*.

The Transformer model architecture consists of a series of encoder blocks interleaving two functions: a multiheaded self-attention computing position-position interactions across the sequence, and a feed-forward network applied independently at each position.

The attention unit:

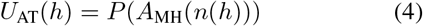

Applies one step of multi-headed scaled dot-product attention to the normalized input, denoted by *n*(*x*), projecting the result into the residual path.

The feed-forward network (with the output state of *P*_1_ defining the “MLP dimension”):

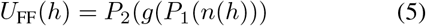

Passes the normalized input through a position-independent multi-layered perceptron (MLP) with activation function *g*(*x*).

The full Transformer block:

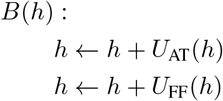

Successively applies the self-attention unit, and the feed-forward network on a residual path.

The Transformer model:

Transformer(*x*) :

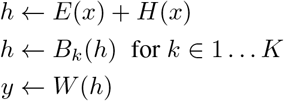

Consists of an embedding step with token *E*(*x*) and positional *H*(*x*) embeddings, followed by *K* layers of Transformer blocks, before a projection *W* to log probabilities. The raw input sequence is represented as a sequence of 1-hot vectors of dimension 25, which is passed through *E*(*x*) the learned embedding layer before being presented to the first Transformer layer.

The models trained in the paper use pre-activation blocks (He et al., 2016), where the layer normalization (Ba et al., 2016) is applied prior to the activation as in Radford et al. (2019b), enabling stable training of deep Transformer networks. No dropout is used. All projections include biases, except for the token and positional embeddings. We use learned token embeddings, and harmonic positional embeddings as in (Vaswani et al., 2017b). The feed-forward network uses the Gaussian error linear unit (Hendrycks & Gimpel, 2016) activation function. We initialize all layers from a zero centered normal distribution with standard deviation 0.02, and re-scale the initialization of the projections into the residual path by 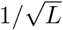 where *L* is the number of residual layers. All biases are initialized to zero. The query, key, and value projections are to *d* dimensions, and the hidden dimension of the feed-forward network is 4*d*.

### 4. Pre-trained Transformer Models

#### UniParc development models

We experimented with Transformer models of various depths, including a 36-layer Transformer with 708.6 million parameters, and a 12-layer model with 85.1M parameters trained. Development models were trained on UniParc. Details are in Table A.1.

#### UniRef models

We train 34-layer models with 669.2M parameters across different datasets and fractions of training data. Additionally we train 6 and 12-layer models. These models are detailed in Table A.2.

#### ESM-1b

The ESM-1b hyperparameter sweep and model is described in detail in Appendix B. In brief, ESM-1b is the result of an extensive hyperparameter sweep that was performed on smaller 12-layer models. ESM-1b is the result of scaling up that model to 33 layers. Compared to the Uniref models, the main changes in ESM-1b are: higher learning rate; dropout after word embedding; learned positional embeddings; final layer norm before the output; and tied input/output word embedding.

#### Pre-training task

The masked language modeling pretraining task follows Devlin et al. (2018b). Specifically, we select as supervision 15% of tokens randomly sampled from the sequence. For those 15% of tokens, we change the input token to a special “masking” token with 80% probability, a randomly-chosen alternate amino acid token with 10% probability, and the original input token (i.e. no change) with 10% probability. We take the loss to be the whole batch average cross entropy loss between the model’s predictions and the true token for these 15% of amino acid tokens. In contrast to Devlin et al. (2018b), we do not use any additional auxiliary prediction losses. The ESM-1b models, as well as the UniParc development models used in visualizations and in the supplemental results are trained with the masking procedure above. The UniRef models used across the experiments of the main text are trained similarly, except that for the 15% of tokens selected as prediction targets, all are replaced by the mask token.

#### Pre-training details

Our model was pre-trained using a context size of 1024 tokens. As most Uniparc sequences (96.7%) contain fewer than 1024 amino acids, the Transformer is able to model the entire context in a single model pass. For those sequences that are longer than 1024 tokens, we sampled a random crop of 1024 tokens during each training epoch. The model was optimized using Adam (*β*_1_ = 0.9, *β*_2_ = 0.999) with learning rate 10^*−*4^. We trained with 131,072 tokens per batch (128 gpus x 1024 tokens). The models follow a warm-up period of 16000 updates, during which the learning rate increases linearly. Afterwards, the learning rate follows an inverse square root decay schedule. All models were trained using the fairseq toolkit (Ott et al., 2019) on 128 NVIDIA V100 GPUs.

### 5. Evaluating the models for downstream tasks

After pre-training the model with unsupervised learning, we can adapt the parameters to supervised tasks. By passing the input sequence (*x*_1_, …, *x*_*n*_) through our pre-trained model, we obtain a final vector representation of the input sequence (*h*_1_, …, *h*_*n*_). During pre-training, this representation is projected to log probabilities (*y*_1_, …, *y*_*n*_). Recall that a soft-max over *y*_*i*_ represents the model’s posterior for the amino acid at position *i*. These final representations (*h*_1_, …, *h*_*n*_) are used directly, or fine-tuned in a task-dependent way by adding additional layers to the model and allowing the gradients to backpropagate through the weights of the pre-trained model to adapt them to the new task. Hidden representations from intermediate layers rather than the final layer can also be used.

### 6. Language Modeling Baselines

In addition to comparing to past work, we also implemented deep learning baselines for our experiments.

#### Frequency (n-gram) models

To establish a meaningful performance baseline on the sequence modeling task (Section 3), we construct n-gram frequency-based models for context sizes 1 ≤*n* ≤10^4^, applying optimal Laplace smoothing for each context size. The Laplace smoothing hyperparameter in each case was tuned on the validation set. ECE is reported for the best left-conditioning n-gram model.

#### Bidirectional LSTM language models

We trained state-of-the-art LSTM (Hochreiter & Schmidhuber, 1997) language models on the UR50 dataset. We use the ELMo model of Peters et al. (2018) which concatenates two independent autoregressive language models with left-to-right and right-to-left factorization. Unlike standard LSTM language models, the ELMo model receives context in both directions and is therefore comparable to the Transformers we train that also use the whole context of the sequence. We train two models: (i) the small model has approximately 28.4M parameters across 3 layers, with an embedding dimension of 512 and a hidden dimension of 1024; (ii) the large model has approximately 113.4M parameters across 3 layers, with an embedding dimension of 512 and a hidden dimension of 4096. The models are trained with a nominal batch size of 32,768, with truncated backpropagation to 100 tokens, dropout of 0.1, learning rate of 8e-4, using the Adam optimizer with betas of (0.9, 0.999), clip norm 0.1 and warmup of 1500 updates using an inverse square root learning rate schedule. We searched across a range of learning rates and found 8e-4 to be optimal.

**Table A.1.** Hyperparameters for development Transformer models trained on UniParc. Embedding dim is the dimension of the hidden states at the output of each transformer block. MLP Dim refers to the width of hidden layer *P*_1_ in the Transformer′s MLPs.

| # Layers | # Heads | Embedding Dim | MLP Dim | # Params | Steps |
| --- | --- | --- | --- | --- | --- |
| 12 | 12 | 768 | 3072 | 85.1M | 1.6M |
| 24 | 12 | 768 | 3072 | 170.2M | 300k |
| 36 | 20 | 1280 | 5120 | 708.6M | 300k |

**Table A.2.** Hyperparameters for UniRef Transformer models. Note: UR100 model stopped making significant progress on valid loss and was stopped at 275K updates.

| # Layers | # Heads | Embedding Dim | MLP Dim | # Params | Data | Steps |
| --- | --- | --- | --- | --- | --- | --- |
| 6 | 12 | 768 | 3072 | 42.6M | UR50/S | 840K |
| 12 | 12 | 768 | 3072 | 85.1M | UR50/S | 840K |
| 34 | 20 | 1280 | 5120 | 669.2M | UR100 | 275K |
| 34 | 20 | 1280 | 5120 | 669.2M | UR50/S | 840K |
| 34 | 20 | 1280 | 5120 | 669.2M | UR50/D | 906K |

### 7. Metric structure experiments

#### Dataset

An orthologous group dataset was constructed from eggNOG 5.0 (Huerta-Cepas et al., 2018) by selecting 25 COG orthologous groups toward maximizing the size of the intersected set of species within each orthologous group. Through a greedy algorithm, we selected 25 COG groups with an intersecting set of 2,609 species. We shrank the dataset above by selecting only one species from each of 24 phyla in order to ensure species-level diversity.

### 8. Remote Homology

#### Dataset

We used the database of SCOP 2.07e filtered to 40% sequence similarity, provided by the ASTRAL compendium (Fox et al., 2014). Following standard practices (Söding & Remmert, 2011), we exclude folds that are known to be related, specifically Rossman-like folds (c.2-c.5, c.27 and 28, c.30 and 31) and four-to eight-bladed *β*-propellers (b.66-b.70). This yields 256,806 pairs of remote homologs at the fold level and 92,944 at the superfamily level, consisting of 217 unique folds and 366 unique superfamilies. We then perform an 80-20 split, and tune our hyperparameters on the “training set” and report results on the held out 20% of the data.

#### Metrics

Given a protein sequence *x*, with final hidden representation (*h*_1_, …, *h*_*n*_), we define the embedding of the sequence to be a vector *e* which is the average of the hidden representations across the positions in the sequence:

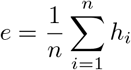

We can compare the similarity of two protein sequences, *x* and *x ′* having embeddings *e* and *e′* using a metric in the embedding space.

We evaluate the L2 distance ‖*e* −*e*^*′*^ *‖*_2_ and the cosine distance *e e*^*′*^*/ e‖ ‖e*^*′*^ *‖*. Additionally we evaluated the L2 distance after projecting the *e* vectors to the unit sphere.

#### Evaluation

To evaluate HHblits (Remmert et al., 2011), first we construct HMM profiles for each sequence using default parameters for ‘hhblits’, except we use 3 iterations. Then, we do an all-to-all alignment using ‘hhalign’ with default parameters, and use the resulting E-value as a measure of similarity. Given a query sequence, a sequence is more similar with a smaller E-value.

The two metrics reported are Hit-10 as introduced in Ma et al. (2014) and AUC. For both metrics, for each sequence, we treat it as a query and we rank each other sequence according to the distance metric used. Following Ma et al. (2014), for evaluation at the fold level, any domain with the same fold is a positive; any domain with a different fold is a negative; and domains belonging to the same superfamily are excluded. For evaluation at the superfamily level, any domain with the same superfamily is a positive; any domain with a different superfamily is a negative; and domains belonging to the same family are excluded. This ensures we specifically measure how well our models do on finding *remote* homologs.

For Hit-10, we consider the query a success if any of the top 10 sequences is a remote homolog. We report the proportion of successes averaged across all queries. For AUC, we first compute the AUC under the ROC curve when classifying sequences by vector similarity to the query. Then, we average the AUC across all query sequences.

We found that cosine similarity results in the best Hit-10 scores, while the L2 with unnormalized vectors result in the best AUC scores, so we report this in Table 2.

#### Implementation

We used the FAISS similarity search engine (Johnson et al., 2017).

## 9. Representational similarity-based alignment of sequences within MSA families

### Family selection

We use the Pfam database (Bateman et al., 2013). We first filtered out any families whose longest sequence is less than 32 residues or greater than 1024 residues in length. We then ranked the families by the number of sequences contained in each family and selected the 128 largest families and associated MSAs. Finally, we reduced the size of each family to 128 sequences by uniform random sampling.

### Aligned pair distribution

For each family, we construct an empirical distribution of aligned residue pairs by enumerating all pairs of positions and indices that are aligned within the MSA and uniformly sampling 50,000 pairs.

### Unaligned pair distribution

We also construct for each family a background empirical distribution of unaligned residue pairs. This background distribution needs to control for within-sequence position, since the residues of two sequences that have been aligned in an MSA are likely to occupy similar positions within their respective unaligned source sequences. Without controlling for this bias, a difference in the distributions of aligned and unaligned pairs could arise from representations encoding positional information rather than actual context. We control for this effect by sampling from the unaligned-pair distribution in proportion to the observed positional differences from the aligned-pair distribution. Specifically, the following process is repeated for each pair in the empirical aligned distribution:

1. Calculate the absolute value of the difference of each residue’s within-sequence positions in the aligned pair.
2. Select a pair of sequences at random.
3. For that pair of sequences, select a pair of residues at random whose absolute value of positional difference equals the one calculated above.
4. Verify that the residues are unaligned in the MSA; if so, add the pair to the empirical background distribution.
5. Otherwise, return to step 2.

This procedure suffices to compute a empirical background distribution of 50,000 unaligned residue pairs.

### Similarity distributions

Finally, for each family and each distribution, we apply the cosine similarity operator to each pair of residues to obtain the per-family aligned and unaligned distribution of representational cosine similarities.

## 10. Linear projections

### Dataset

We construct five-fold cross validation datasets with structural hold-outs at the family, superfamily, and fold level using SCOPe 2.07 (Fox et al., 2014). We use the full version of SCOPe 2.07, clustered at 90% sequence identity, generated on January 23, 2020, and extract the domain annotations with labels. There are 19,695 domains. Then, independently for each hold-out level, we split the domains at the hold-out level into 5 equal sets, i.e. five sets of folds, superfamilies, or families. This ensures that for each partition, structures having the same classification do not appear in both the train and test sets. For a given classification level each structure appears in a test set once, so that in the cross validation experiment each of the structures will be evaluated once. Scores reported are the mean and standard deviation over each of the five test sets.

For each domain, we first obtain the sequence *x* whose residues align with the domain specification. To construct the secondary structure labels, we take each CIF file (pulled from PDB), and run DSSP (Joosten et al., 2010; Kabsch & Sander, 1983). If *x* has any residues where DSSP has not provided a secondary structure label, we mark them as missing data and do not supervise for those positions.

To construct the contact map, we obtain C_*β*_ coordinates from the structure portion of the CIF file (C_*α*_ in the case of glycine), defaulting to NaN where information is missing, and finally calculating pairwise distances and thresholding at 8Å. Similar to secondary structure, we do not supervise over NaNs.

We discard any domains where (1) DSSP fails, (2) we are unable to align the sequence to the structure, or (3) the domain is longer than 1023 residues. This leaves 15,297 domains.

For the MSA baselines, we query each sequence against the Uniclust30, 2017 database (Mirdita et al., 2017) with HH-blits (Remmert et al., 2011) using the default settings with additional parameters (n=3, 1e-3). For secondary structure prediction, we construct HMM profiles using HHmake (default settings). For contact prediction, we apply CCMpred (Seemayer et al., 2014) implementation of pseudolikelihood maximization (Balakrishnan et al., 2011; Ekeberg et al., 2013) using default settings with (n=500) to each MSA, from which we extract both an output matrix (ℝ^*L×L*^), as well as a sequence profile (ℝ^*L×K*^) where *L* is the length of the sequence and *K* is the size of the amino acid vocab, i.e. 25.

#### Representations

To obtain sequence representations, we provide the sequence of the domain as input to a forward pass of the Transformer model. We retrieve the activations into the final multi-head attention (after the layer normalization), using this as the matrix of sequence representations (ℝ^*L×d*^) where *d* is the hidden dimension of the model.

#### Secondary structure projections

For secondary structure, we fit a multi-class logistic regression taking as input an individual representation *h*_*i*_ and as output the secondary structure label from DSSP. We observe that the logistic regression model’s performance does not change with penalty settings (L1, L2, no penalty); therefore we report the result where the L2 penalty is applied during training.

#### Contact projections

For contacts, we fit two linear projections. Given two representations *h*_*i*_, *h*_*j*_ at positions *i* and *j*, we regress to whether residues at positions *i* and *j* are in contact:

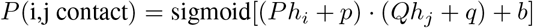

With learned projections *P, Q*, vector biases *p, q*, and scalar bias *b*. We use the AdamW optimizer (Loshchilov & Hutter, 2017) to fit the projections and bias term.

For each partition we set aside approximately 12.5% of the training set as validation. We sweep over a range of projection dimensions (32 to 512), learning rates (1e-6 to 1e-2) and weight decay values (0 to 0.9). Based on the best validation Top-L, long-range precision score, we set the projection dimension to 128, the learning rate to 1e-4, and the optimizer weight decay to 0.2. We observe that the precision score does not improve with an increased projection dimension over 128.

The above setup for contact projections only applies to the Transformer models and the sequence profile baseline. No supervision is applied to the CCMpred output.

### 11. Single-family data and analysis

For each of the three domains used, we extracted all domain sequences from the Pfam dataset (Bateman et al., 2013) and located the subset of PDB files containing the domain, using the latter to derive ground truth secondary structure labels (Kabsch & Sander, 1983).

Pre-training is with the masked language modeling objective, using the same hyperparameters as used to train the UniParc development models. The domain sequences were randomly partitioned into training, validation, and testing datasets. For each family, the training dataset comprises 65,536 sequences, the validation dataset comprises either 16,384 sequences (PF00005 and PF00069) or 8,192 sequences (PF00072), and the test dataset comprises the remainder.

Each Pfam family also forms an evaluation dataset for linear projection; from the sequences with corresponding crystal structures, the training dataset comprises 80 sequences and the test dataset comprises the remainder.

### 12. Secondary structure prediction. Deep neural networks and feature combination

We use features from the final hidden representations of the models. We removed the final embedding layer, added layer norm, and applied a top-level architecture following (Klausen et al., 2019). In particular, this top-level architecture consists of two parallel convolution layers and an identity layer, whose outputs are concatenated in the feature dimension and fed to a two layer bidirectional LSTM containing 1024 hidden units and dropout *p* = 0.5. The output is then projected to an 8-dimensional feature vector at each position and the model is trained with a categorical cross-entropy loss with the Q8 labels. The training data was obtained from the (Klausen et al., 2019). Secondary structure labels for the CASP13 test set were constructed using DSSP.

In feature combination experiments, we used the features provided by (Klausen et al., 2019) which were generated using MMseqs2 on the Uniclust90 dataset released April 2017. For CASP13 experiments, we generated these features using code provided by (Klausen et al., 2019) on CASP13 domains.

As a baseline, we reimplemented (Klausen et al., 2019) by replacing the Transformer features with the MMseqs2 features and keeping the top-level architecture. For feature combination experiments, we projected (a) the features from this baseline and (b) the features from the Transformer to the same dimension (512 units), concatenated along the feature dimension, and fed the resulting tensor to a two layer bidirectional LSTM with 512 hidden units and dropout *p* = 0.3.

To check our dataset construction, we used the pretrained weights provided by (Klausen et al., 2019) and evaluated their model directly in our evaluation pipeline. We were able to reproduce the values reported in (Klausen et al., 2019).

### 13. Contact prediction. Deep neural networks and feature combination

#### Data

We use the datasets and features distributed with Wang et al. (2017) and Xu (2018). The base features are those used by RaptorX (Xu, 2018) a state-of-the-art method in CASP13, including sequence features, PSSM, 3-state secondary structure prediction, predicted accessibility, one-hot embedding of sequence, and pairwise features APC-corrected Potts model couplings, mutual information, pair-wise contact potential.

We use the training, standard test set, and CASP11 test set from Wang et al. (2017). We use the CASP12 test set from Xu (2018). For the CASP13 test set we use the 34 publicly released domains.

Wang et al. (2017) established training and test sets as follows. The train (6,367 proteins), valid (400 proteins) and test (500 proteins) datasets were selected as subsets of PDB25 (each protein having <25% sequence similarity). Proteins having sequence similarity >25% or BLAST E-value <0.1 with any test or CASP11 protein were excluded from training data.

All our MSAs (used for the avg and cov combination methods) are constructed by running HHblits (Remmert et al., 2011) with 3 iterations and E-value 0.001 against Uniprot20 released on 2016-02; except for CASP12 and CASP13 where we used the four different MSAs released with and described in Xu (2018). Note that for the Transformer pretraining UniRef50 from 2018-03 was used; hence no data which was not already available prior to the start of CASP13 was present during either pre-training or contact prediction training.

#### Model architecture

On top of the sequence and pairwise features we use a depth-32 residual network (ResNet) model to predict binary contacts. The ResNet model architecture is similar to Wang et al. (2017) and Xu (2018).

The first component of the ResNet is a learned sequence pipeline 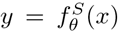 which maps sequence features *x* ∈ 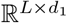 to *y* ∈ 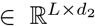 with *L* the length of the protein. Though 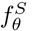 could be a 1D convolutional network or residual network as in Wang et al. (2017), we chose our sequence net to be a simple linear projection from input dimension *d*_1_ to *d*_2_ = 128 dimensions. The input dimension *d*_1_ is either 46 (RaptorX only), 1280 (Transformer hidden state), or 1326 (feature combination). We varied *d*_2_ and empirically determined 128 to be optimal.

The 128-D output *y* of the sequence net gets converted to pairwise matrix features *z*_1_ with 256 feature maps, by an outer concatenation operation; i.e. at position *i, j* we concatenate *y*_*i*_ and *y*_*j*_ along the feature dimension, giving rise to 2 × *d*_2_ feature maps. This 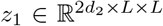 is then concatenated in the first (feature map or channel) dimension, with the pairwise features *z*_2_ ∈ ℝ^6*×L×L*^ i.e. the pairwise RaptorX features described in previous subsection and/or the msa embedding covariance features (*z*_3_ ∈ R^256*×L×L*^) described in the next subsection. As such the concatenated *z* ∈ ℝ262*×L×L* or *z* ∈ ℝ ^518*×L×L*^.

The final component is the actual 2D residual network operating in *z*, which computes the binary contact probability 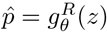 with 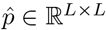 and 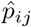 the continuous predicted probability of position *i* and *j* of the protein being in contact. The ResNet has an initial 1 ×1 convolutional layer going to *d*_3_ = 128 feature maps, followed by MaxOut over the feature maps with stride 2, reducing to 64 feature maps. After this, there are 32 residual blocks. Each residual block has on its weight path consecutively BatchNorm – ReLU - Conv 3 × 3 (64 feature maps) - Dropout (0.3) - ReLU - Conv 3 × 3 (64 feature maps). The residual blocks have consecutive dilation rates of 1,2,4. This follows Adhikari (2019). The final output is computed with a Conv 3 ×3 (1 output feature map) and sigmoid to produce probability of contact 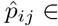[0,1]. As such there are 66 convolutional layers in the main 2D ResNet.

Note that a number of shortcomings exist from our pipeline to CASP13 winners (Senior et al., 2018; Xu, 2018); most importantly we use an earlier training dataset of PDB structures compiled from PDB dated Feb 2015 by Wang et al. (2017), additionally we do not incorporate more recent developments like distance distribution prediction, sliding window on small crops allowing deeper ResNets, auxiliary losses like torsion angles, or data augmentation.

For reference, the officially released AlphaFold (Senior et al., 2018) predictions achieve a top-L/5,LR and top-L,LR precision on the same subset of CASP-13 targets of 75.2% and 52.2% respectively. The discrepancies in the pipeline explain why our best precisions using RaptorX features are about 7-9% lower (compare CASP13-AVG (a): 68.0% / 43.4%)

#### MSA Embedding feature combination

We construct features based on the embedding of the MSA of a protein sequence in our training data. We denote the original protein in our labeled dataset, i.e. query sequence *x* of length *L*, to have corresponding embedding *h* = Transformer(*x*) ∈ ℝ^*L×d*^, and the embedding of the *i*-th position to be *h*_*i*_ ∈ ℝ^*d*^. Typically *h* is the last hidden state from the pre-trained Transformer model. The *m*th sequence in the MSA is *x*^*m*^, with corresponding embedding *h*^*m*^. *m* ∈ [0, *M* [with *M* the MSA depth. The embeddings are computed by embedding the original sequence *x*^*m*^ without inserting gaps (there is no gap character in our vocabulary), then realigning the embedding according to the alignment between *x*^*m*^ and query sequence *x* by inserting 0-vectors at position *i* if the 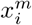 is the gap character; ie 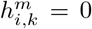. We also use indicator variable 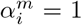 if 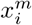 is non-gap (match state), or 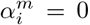 if 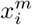 is gap. We further compute sequence weights *w*^*m*^ as the commonly used debiasing heuristic to reduce the influence of the oversampling of many similar sequences. The weights are defined in the usual way with 70% sequence similarity threshold: sequence weight 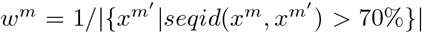 which is the inverse of the number of sequences *x*^*m*^ that are more than 70% similar to the sequence *x*^*m*^ i.e. hamming distance less than 0.3L.

Now we introduce the average embedding over an MSA:

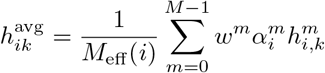

with per-position denominator 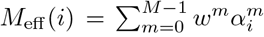 This is effectively a weighted average over the sequence embeddings in the MSA. Note that if the embeddings were one-hot encodings of AA identities, we would recover the position probability matrix (except the absence of a pseudo-count).

Similarly; we introduce the (uncentered) covariance of the embeddings, with PCA-projected 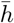

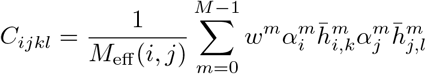

With pairwise position denominator 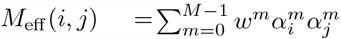.

Note that to make above covariance embedding feasible, we first reduce the dimensionality of the embeddings by projecting onto the first 16 PCA directions: 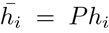 with *P* ∈ ℝ ^16*×d*^, giving rise to a covariance per pair of positions *i, j* and pair of interacting PCA components *k, l* of *L* × *L* ×16 ×16. The 256 different *k, l* pairs of *C*_*ijkl*_ ∈ ℝ^*L×L×*16*×*16^ will now become the feature maps of *z* ∈ R^256*×L×L*^, such that *z*_16*k*+*l,i,j*_ = *C*_*ijkl*_. We tried training (rather than fixed PCA) the projection of the features 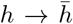 before covariance (learned linear projection *P* or training a 3-layer MLP). We also varied the formulation to center the embeddings over the MSA (normal covariance) and to rescale the feature maps with a pre-computed mean and standard deviation for each feature map corresponding to a pair of *k, l*. We found no gains from these variations over the current formulation. Note that centering with the average *h*^avg^ as in normal empirical covariance calculation, introduces a shift that is independent per protein (because specific to the MSA), and independent per position. Therefore it is not unexpected that the uncentered covariance gives better (more consistent) features.

### 14. Mutational Effect

#### Datasets

We used two datasets of variant effect measurements compiled by Gray et al. (2018) and Riesselman et al. (2018). The first dataset is a collection of 21,026 measurements from nine experimental deep mutational scans. The second dataset contains 712,218 mutations across 42 deep mutational scans.

#### Fine-tuning procedure

To fine-tune the model to predict the effect of changing a single amino acid or combination of amino acids we regress the scaled mutational effect with:

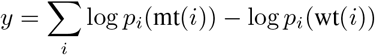

Where mt(*i*) is the mutated amino acid at position *i*, and wt(*i*) is the wildtype amino acid. The sum runs over the indices of the mutated positions. As an evaluation metric, we report the Spearman *ρ* between the model’s predictions and experimentally measured values.

### 15. Area under the ROC curve

For a binary classification task, the ROC curve plots the true positive rate against the false positive rate at various classification thresholds. The area under the ROC curve gives a measure that quantifies the model’s ability to distinguish between classes. Intuitively a perfect classifier has an AUC of 1, while a uniform random classifier has an AUC of 0.5.

## B. ESM-1b Hyperparameter Optimization

### Experimental setup

We perform a systematic analysis on Transformer models with 100M parameters. We train models on the UniRef50 dataset following the same methodology described in the rest of this work.

After identifying the best performing settings on 100M parameter models, we explore scale by training 650M parameter models. All models are trained with a target batch size of 1M tokens. To accommodate the large batch size, we use gradient accumulation and distributed data parallel. Under this setup, each epoch of a 100M parameter model completes in 1.8 hours on 32 GPUs. Each epoch of a 650M parameter model completes in 8.5 hours on 64 GPUs.

When studying architectural variants, we assess the quality of representations from each model after 10 or 12 epochs of pre-training. We observed that relative performance ranking of the models does not change after this point. Notably, this is still early in training; our best performing model, ESM-1b, is trained for 56 epochs.

#### Hyperparameter: Masking and data setup

Protein language models are trained with a masked language modeling objective, wherein each input sequence is corrupted by replacing a fraction of the tokens with a special mask token. We train 100M parameter models for 10 epochs and compare their performance on the CB513 test set. We investigate four masking strategies:

- *All masks*: following supplemental section 4, 15% of the input tokens are replaced with a mask token and predicted.
- *All random (uniform)*: 15% of the input tokens are replaced with an amino acid selected uniform randomly and predicted.
- *All random (frequency)*: 15% of the input tokens are replaced with an amino acid selected according to their frequency in the dataset and predicted.
- *BERT*: 15% of the input tokens are selected and predicted. Of these, 80% are replaced with mask token; 10% with a uniform random amino acid; 10% not changed.

In all cases, we follow Liu et al. (2019) and dynamically mask the sequences, such that a new mask is randomly selected at each epoch. Since the input data changes across runs, language modeling perplexities cannot be fairly compared. Therefore, we evaluate the downstream performance of the models on a secondary structure benchmark. Table .**B. 1** finds that the *BERT* masking pattern performs better than the other masking patterns and is therefore used for all model variations below.

**Table B.1.** Comparison of masking patterns, 8-class secondary structure prediction accuracy. Models are pre-trained for 10 epochs on Uniref50. The model and all other hyperparameters remain fixed between experiments. The BERT masking pattern performs best and is used for future experiments.

| Masking pattern | CB513 |
| --- | --- |
| All masks | 60.4 |
| All random (uniform) | 59.3 |
| All random (frequency) | 59.0 |
| BERT | 60.8 |

#### Hyperparameter: Dynamic batching

Models in section 4 were trained with dynamic batching, which results in a single sequence in each sample in the batch. This design choice contrasts with existing language modeling works. For example, in NLP, Liu et al. (2019) and Devlin et al. (2018a), use a static batching approach, wherein multiple proteins are concatenated in the same batch along the sequence dimension. This approach is common in NLP, as sentences that are nearby in a corpus generally relate to the same topic. As this situation is not the case in protein language modeling, we analyze the impact of static batching schemes in protein language models, finding that they reduce model performance (Table B.2). 100M parameter models are evaluated on the secondary structure downstream task after 10 epochs.

**Table B.2.** Comparison of batching modes, 8-class secondary structure accuracy. Batching modes are investigated on 100M parameter models and trained for 10 epochs. All models are trained with a context size of 1024 tokens. In the static batching mode, sequences longer than 1024 can span multiple batches. We crop long sequences for the other batching modes considered. The dynamic batching scheme we used performs significantly better than static batching modes.

| Batching mode | CB513 |
| --- | --- |
| Static | 56.2 |
| Static [cropping long sequences] | 57.0 |
| Dynamic | 60.8 |

#### Hyperparameters: further sweeps

After fixing the data distribution to use the BERT masking scheme with dynamic batching, we next perform a hyperparameter sweep to identify the best learning rates, initializations and layer norm placement. We also propose a token dropout scheme which further improves performance on downstream tasks.

#### Initializations

We compare our initializations to the initializations presented in Liu et al. (2019), finding that they perform similarly (Table B.3).

**Table B.3.** Comparison of initializations, 8-class secondary structure accuracy. 100M parameter models are trained for 10 epochs and evaluated on CB513.

| Initializations | CB513 |
| --- | --- |
| Radford et al. (2019a) | 60.9 |
| Liu et al. (2019) | 60.8 |

#### Layer norm placement

Recent works Child et al. (2019) and Shoeybi et al. (2020) have suggested that pre-activation layer norm results in more stable training for larger models. We investigate the impact of this choice in a smaller controlled setting using 100M parameter transformer models, finding that pre-activation layer norm improves performance on downstream tasks (Table B.4). To account for the lack of a final layer norm, we additionally add a final layer norm before the linear output projection, improving performance (Table B.5).

**Table B.4.** Comparison of layer norm placement, 8 class secondary structure accuracy. 100M parameter models are trained for 10 epochs and evaluated on CB513.

| Model | CB513 |
| --- | --- |
| Post-activation | 60.8 |
| Pre-activation | 61.1 |

**Table B.5.** Including a final layer norm before the language modeling head performs best, 8-class secondary structure accuracy. 100M parameter models are trained for 12 epochs.

| Final layer norm | CB513 |
| --- | --- |
| No | 61.3 |
| Yes | 61.9 |

#### Token dropout

Usually, masked language models are pretrained with corrupted inputs and fine-tuned with complete sequences. We hypothesize that this shift in data distribution reduces performance on downstream tasks. Therefore, we propose a token dropout scheme which replaces the mask token embedding with a fixed tensor of zeros. As 10 −15 of positions are masked, the zero tensors cause a change in the mean statistics of the word embeddings. We therefore adjust the distribution during fine-tuning by multiplying the word embeddings by a fixed constant. Formally, if a mask token is introduced during pre-training with probability *p* = 0.15 · 0.8, then during fine-tuning, we multiply the embeddings by 1*/p*. We find that this significantly improves performance (Table B.6).

**Table B.6.** Token dropout performs better than mask tokens, 8-class secondary structure accuracy. 100M parameter models are trained for 12 epochs.

| Masking | CB513 |
| --- | --- |
| Mask Tokens | 61.9 |
| Token Dropout | 62.6 |

#### Learning rate

Next, we investigate learning rate, finding that a peak learning rate of 4 · 10^*−*4^ performs best (Table B.7). Higher learning rates result in instability, while lower learning rates result in lower performance. In all cases, learning rate is warmed up linearly for 16000 steps and then decayed following an inverse square-root schedule (Raffel et al., 2020).

**Table B.7.** A peak learning rate of 4 · 10^*−*4^ performs best, 8 class secondary structure accuracy. 100M parameter models are trained for 10 epochs.

| Learning rate | CB513 |
| --- | --- |
| $1 \cdot 10^{-4}$ | 61.9 |
| $2 \cdot 10^{-4}$ | 63.7 |
| $4 \cdot 10^{-4}$ | 65.2 |

#### Tying embeddings

Although all experiments above are performed with tied input and output embeddings, we investigate whether learning these separately could improve performance. Our results (Table B.8) indicate that sharing the weights negatively impacts performance. Therefore, we maintain our initial setup.

**Table B.8.**
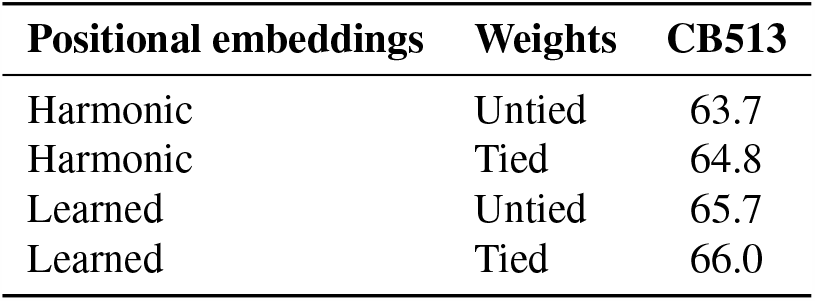
Comparison of tied embeddings and l.earned or harmonic positional embeddings, 8 class secondary structure accuracy. 100M parameter models are trained for 17 epochs.

| Positional embeddings | Weights | CB513 |
| --- | --- | --- |
| Harmonic | Untied | 63.7 |
| Harmonic | Tied | 64.8 |
| Learned | Untied | 65.7 |
| Learned | Tied | 66.0 |

#### Positional embeddings

We also investigate the impact of learning positional embeddings compared to fixed harmonic embeddings (Vaswani et al., 2017a), finding that learned positional embeddings perform better (Table B.8).

**Figure S1.**
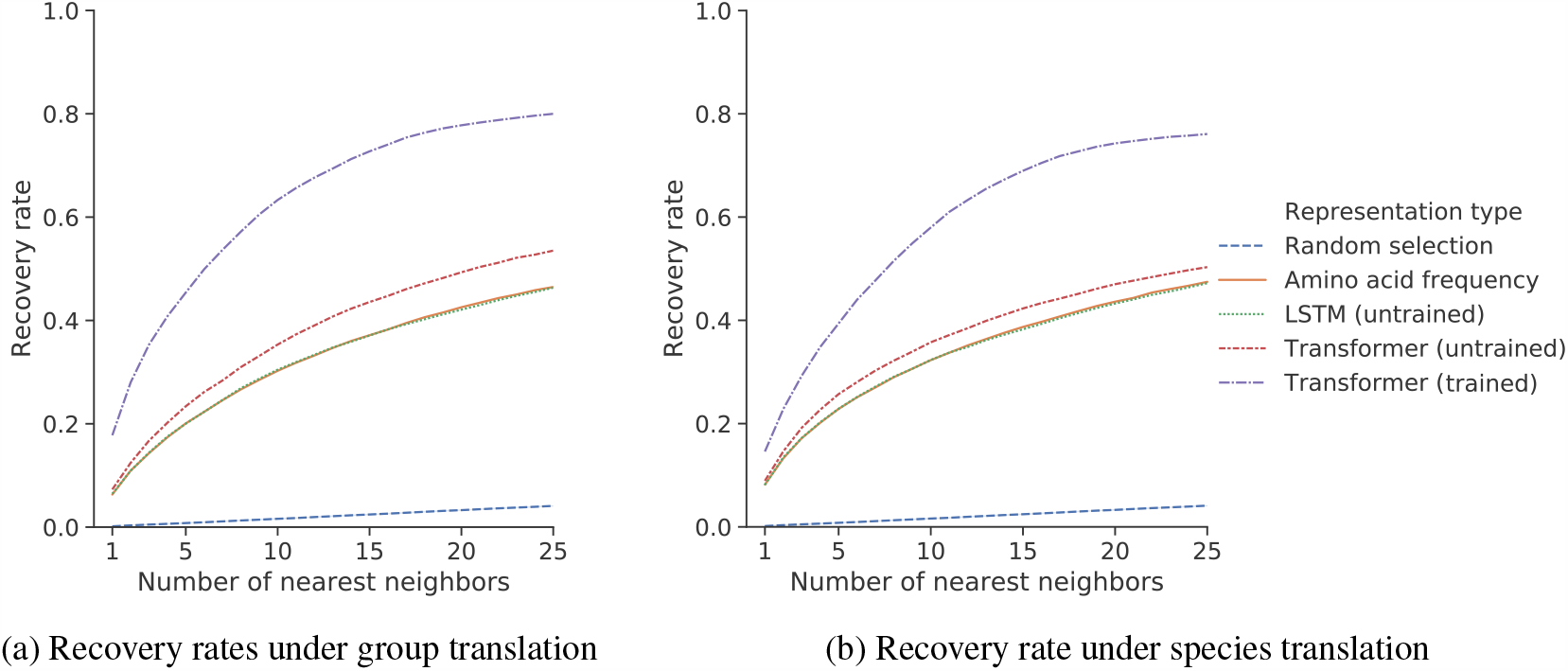
Learned sequence representations can be translated between orthologous groups and species. Depicted is the recovery rate of nearest-neighbor search under (a) orthologous group translation, and (b) species translation. In both settings, the trained Transformer representation space has a higher recovery rate. Results shown for 36-layer dev Transformer pre-trained on UniParc. To define a linear translation between protein *a* and protein *b* of the same species, we define the source and target sets as the average of protein *a* or protein *b* across all 24 diverse species. If representation space linearly encodes orthology, then adding the difference in these averages to protein *a* of some species will recover protein *b* in the same species. We use an analogous approach to translate a protein of a source species *s* to its ortholog in the target species *t*. Here, we consider the average representation of the proteins in *s* and in *t*. If representation space is organized linearly by species, then adding the difference in average representations to a protein in species *s* will recover the corresponding protein in species *t*.

**Table S1.** Area under the ROC curve (AUC) of per-residue representational cosine similarities in distinguishing between aligned and unaligned pairs of residues within a Pfam family. Results displayed are averaged across 128 families.

| Representation type | Overall | Identical amino acid pairs | Distinct amino acid pairs |
| --- | --- | --- | --- |
| Transformer (trained) | **0.841** | **0.870** | **0.792** |
| Transformer (untrained) | 0.656 | 0.588 | 0.468 |

**Table S2.**
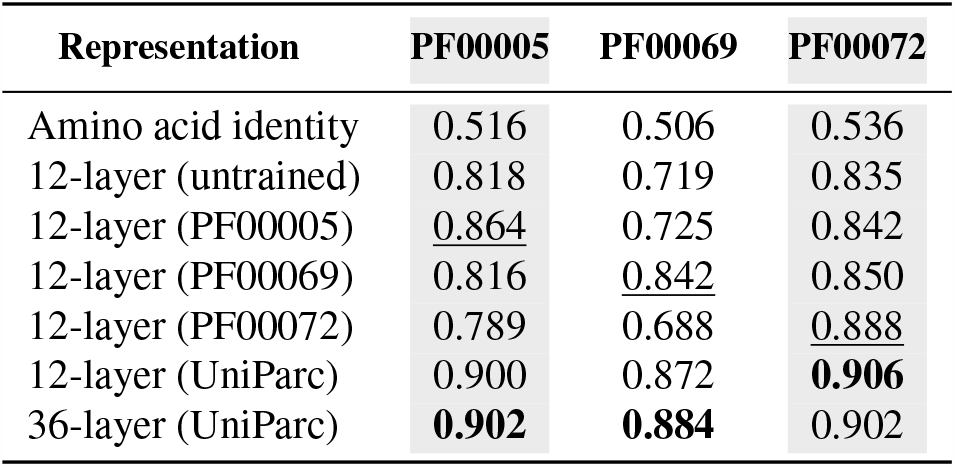
Three-class secondary structure prediction accuracy by linear projection. Learning across many protein families produces better representations than learning from single protein families. Transformer models are trained on three PFAM families: ATP-binding domain of the ABC transporters (PF00005), Protein kinase domain (PF00069), and Response regulator receiver domain (PF00072). The single-family models are contrasted with models trained on the full UniParc data. Comparisons are relative to the family (columnwise), since each of the families differ in difficulty. Underline indicates models trained and evaluated on the same family. Representations learned from single families perform well within the family, but do not generalize as well to sequences outside the family. Representations trained on UniParc outperform the single-family representations in all cases.

| Representation | PF00005 | PF00069 | PF00072 |
| --- | --- | --- | --- |
| Amino acid identity | 0.516 | 0.506 | 0.536 |
| 12-layer (untrained) | 0.818 | 0.719 | 0.835 |
| 12-layer (PF00005) | <u>0.864</u> | 0.725 | 0.842 |
| 12-layer (PF00069) | 0.816 | <u>0.842</u> | 0.850 |
| 12-layer (PF00072) | 0.789 | 0.688 | <u>0.888</u> |
| 12-layer (UniParc) | 0.900 | 0.872 | **0.906** |
| 36-layer (UniParc) | **0.902** | **0.884** | 0.902 |

**Figure S2.**
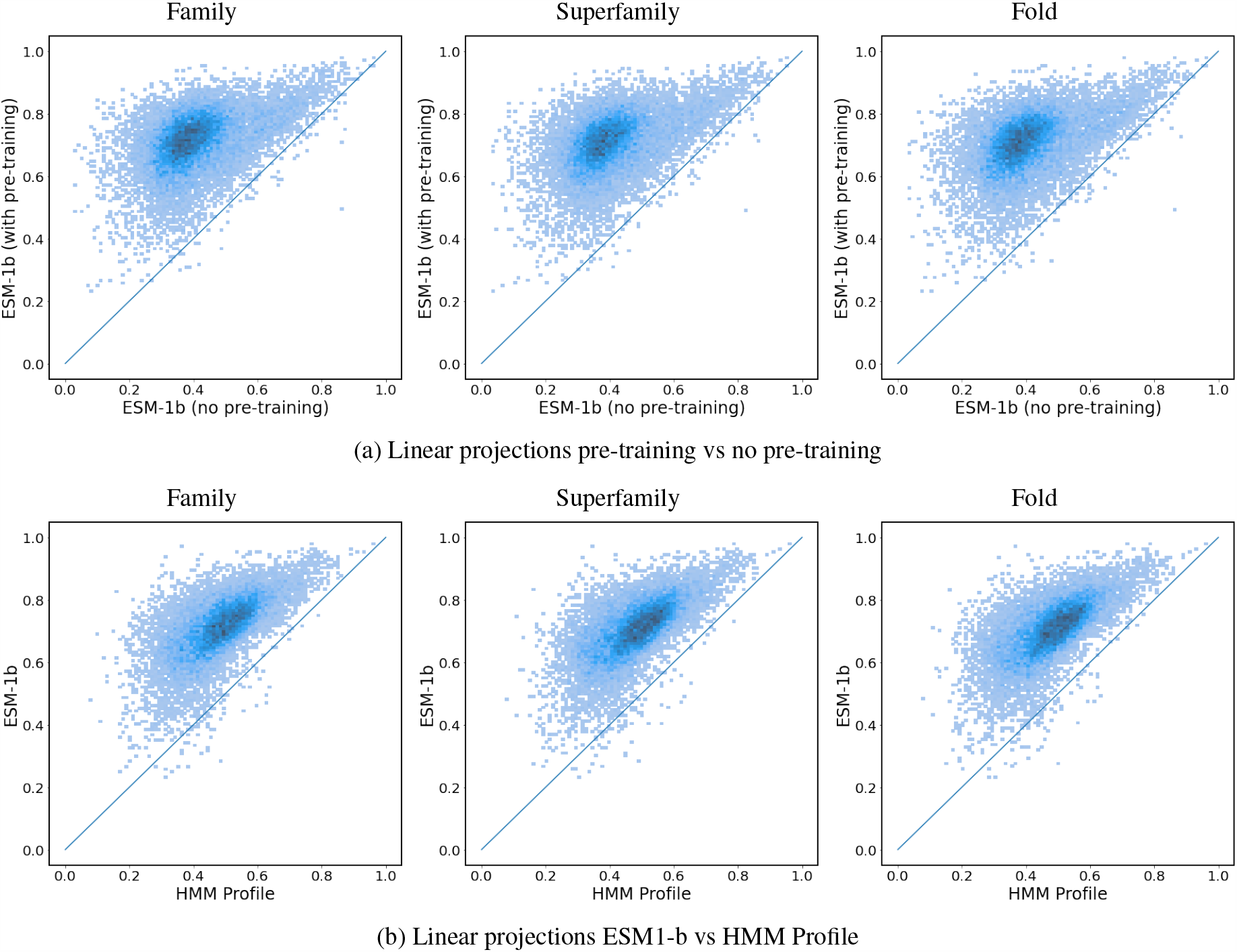
Secondary structure prediction 8-class accuracy distributions for linear projections. (a) Comparison with and without pretraining; (b) comparison of ESM-1b Transformer representations with HMM sequence profiles. Density is indicated by color.

**Figure S3.**
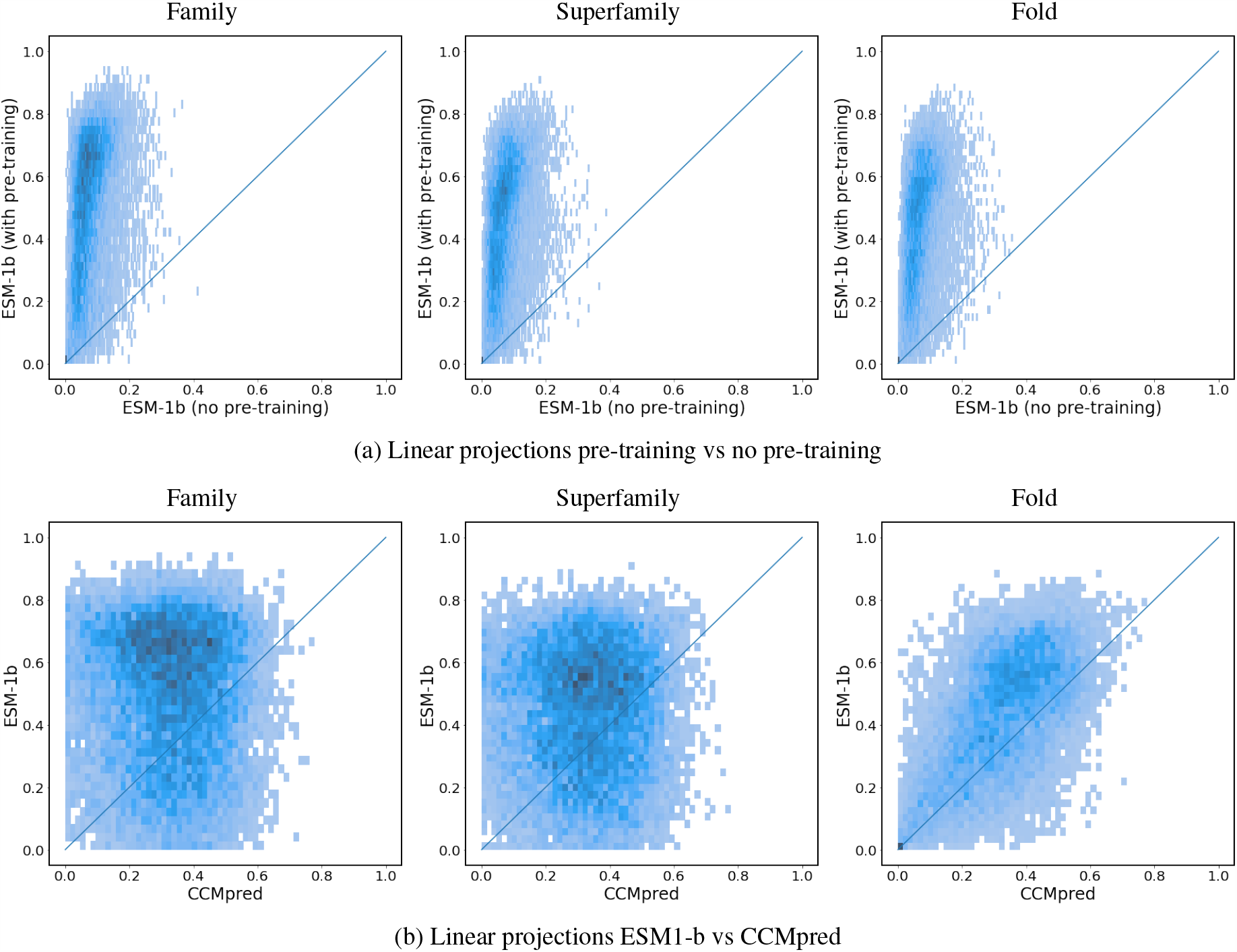
Contact prediction Top-L long-range precision distributions for linear projections. (a) Comparison with and without pretraining; (b) comparison of ESM-1b Transformer representations with CCMpred predictions. Density is indicated by color.

**Figure S4.**
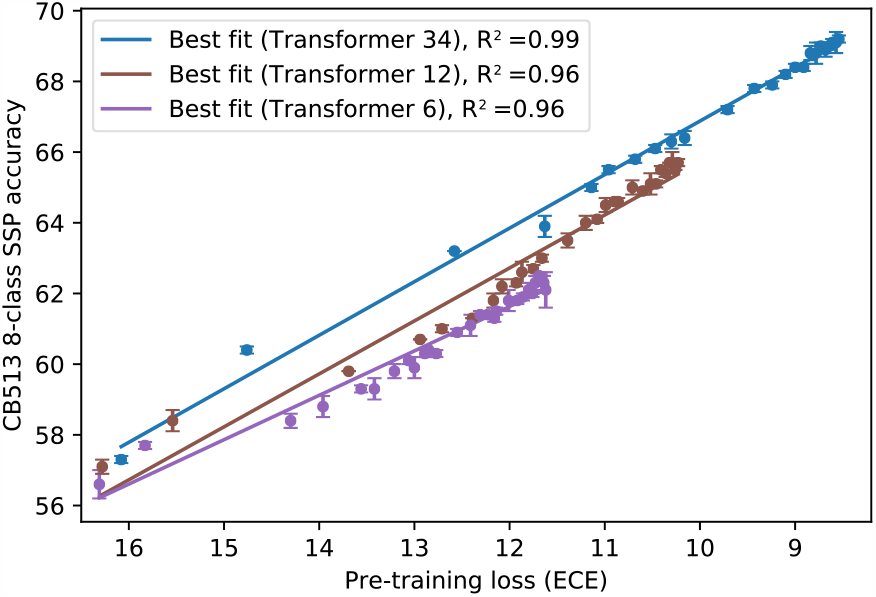
Eight-class secondary structure prediction accuracy as a function of pre-training ECE. A deep secondary structure predictor is trained using features from Transformer models over the course of pre-training on UR50/S. The Netsurf training sequences and CB513 test set are used. Averages across three seeds of the downstream model per pre-training checkpoint are plotted, with line of best fit for each Transformer.

**Table S3.** Additional metrics. Feature combination for supervised contact prediction. AVG corresponds to the average of the metrics over the different MSAs, while in ENS the probabilities are averaged (ensembled) over the different MSA predictions before computing the metrics.

| Metric: top- |  | L/5,LR | L,LR | L/5,MR | L,MR | L/5,SR | L,SR |
| --- | --- | --- | --- | --- | --- | --- | --- |
| Test | (a) RaptorX | 84.3 ± .2 | 59.4 ± .2 | 74.4 ± .1 | 33.0 ± .0 | 71.6 ± .1 | 25.8 ± .0 |
| | (b) +direct | 86.7 ± .3 | 61.7 ± .4 | 76.5 ± .3 | 33.8 ± .1 | **73.6 ± .2** | **26.1 ± .1** |
| | (c) +avg | **87.7 ± .3** | 62.9 ± .4 | 77.4 ± .2 | **34.0 ± .1** | **73.7 ± .3** | **26.1 ± .1** |
|  | (d) +cov | **87.8 ± .3** | **63.3 ± .2** | **77.6 ± .2** | **34.0 ± .1** | **73.7 ± .2** | **26.1 ± .0** |
| CASP11 | (a) RaptorX | 77.5 ± .4 | 53.8 ± .3 | 75.0 ± .5 | 35.6 ± .2 | 72.1 ± .6 | 28.6 ± .2 |
| | (b) +direct | 78.3 ± .1 | 55.0 ± .1 | 76.2 ± .4 | 35.9 ± .2 | 74.0 ± .5 | 28.8 ± .2 |
| | (c) +avg | **80.4 ± .5** | 56.6 ± .4 | **76.5 ± .4** | 36.3 ± .2 | **73.8 ± .5** | 28.8 ± .1 |
|  | (d) +cov | **80.3 ± .3** | **56.8 ± .2** | **76.6 ± .4** | **36.5 ± .2** | **74.0 ± .4** | **29.0 ± .0** |
| CASP12-AVG | (a) RaptorX | 72.7 ± .6 | 51.1 ± .2 | 68.3 ± .5 | 31.2 ± .3 | 66.5 ± .2 | 26.3 ± .1 |
| | (b) +direct | 74.0 ± .8 | 51.5 ± .5 | 70.7 ± .7 | **32.4 ± .3** | 68.9 ± .9 | **27.2 ± .2** |
| | (c) +avg | 74.4 ± 1.4 | 52.4 ± .5 | **71.7 ± .6** | **32.2 ± .3** | **70.1 ± .2** | 26.9 ± .2 |
| | (d) +cov | **76.6 ± .7** | **53.0 ± .3** | 70.1 ± .3 | 31.9 ± .3 | 69.1 ± .5 | 26.6 ± .1 |
| CASP12-ENS | (a) RaptorX | 77.1 ± .9 | 54.5 ± .4 | 70.6 ± .6 | 32.4 ± .4 | 68.6 ± .4 | 27.0 ± .1 |
| | (b) +direct | 77.0 ± .6 | 53.5 ± .6 | **71.9 ± .9** | **33.1 ± .3** | 69.8 ± .7 | **27.6 ± .2** |
| | (c) +avg | 76.7 ± 1.4 | 54.4 ± .7 | **74.1 ± .8** | **33.0 ± .3** | **71.5 ± .3** | 27.4 ± .2 |
| | (d) +cov | **79.7 ± .8** | **55.3 ± .2** | 72.7 ± .5 | 32.7 ± .3 | 71.0 ± .6 | 27.2 ± .2 |
| CASP13-AVG | (a) RaptorX | 68.0 ± .9 | 43.4 ± .4 | 71.3 ± .4 | 36.5 ± .3 | 68.8 ± 1.0 | 28.4 ± .0 |
| | (b) +direct | 67.4 ± .8 | 43.7 ± .4 | 69.5 ± .9 | 35.5 ± .4 | 68.1 ± .4 | 28.3 ± .2 |
| | (c) +avg | 68.1 ± 1.6 | **44.8 ± .8** | **73.0 ± .6** | **36.9 ± .1** | **71.2 ± .6** | **28.6 ± .2** |
| | (d) +cov | **70.3 ± 1.3** | **45.2 ± .5** | **73.5 ± 1.4** | **37.0 ± .2** | 70.2 ± .8 | **28.6 ± .3** |
| CASP13-ENS | (a) RaptorX | **72.1 ± .8** | 46.3 ± .5 | 73.6 ± .4 | **38.1 ± .3** | 71.0 ± 1.4 | **29.6 ± .1** |
| | (b) +direct | 68.0 ± .7 | 45.0 ± .6 | 71.4 ± 1.2 | 36.6 ± .4 | 70.2 ± .2 | 29.1 ± .2 |
| | (c) +avg | 70.8 ± 2.2 | 46.4 ± 1.1 | **75.4 ± .5** | **38.1 ± .2** | **73.6 ± .5** | 29.3 ± .2 |
| | (d) +cov | 71.9 ± 1.9 | **47.2 ± .4** | **75.2 ± 1.5** | **38.3 ± .4** | **72.3 ± .8** | 29.3 ± .2 |

**Figure S5.**
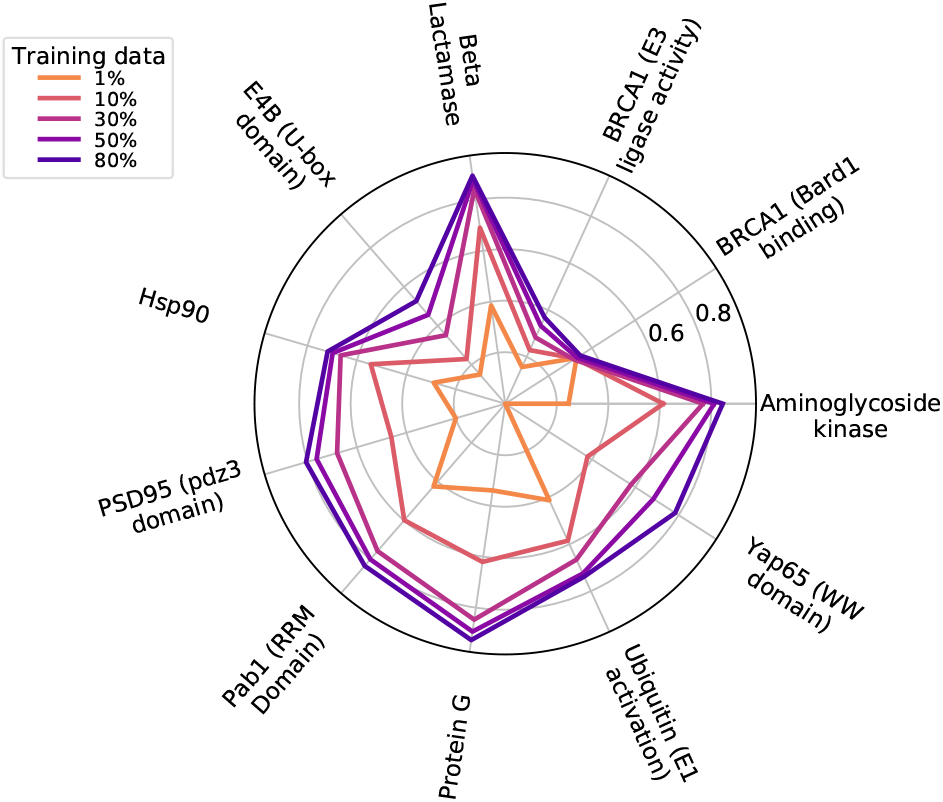
After pre-training, the Transformer can be adapted to predict mutational effects on protein function. The 34-layer Transformer model pre-trained on UR50/S is fine-tuned on mutagenesis data. Spearman *ρ* on each protein when supervised with smaller fractions of the data.

**Table S4.** Aggregate spearman *ρ* measured across models and datasets. Mean and standard deviations of spearman *ρ* performance for the fine-tuned Transformer-34 on intraprotein tasks. Performance was assessed on five random partitions of the validation set. Model pre-trained on UR50/S.

| Amount of training data<br>Protein | 1% data | 10% data | 30% data | 50% data | 80% data |
| --- | --- | --- | --- | --- | --- |
| Aminoglycoside kinase | $0.25 \pm 0.07$ | $0.61 \pm 0.02$ | $0.77 \pm 0.01$ | $0.81 \pm 0.01$ | $0.84 \pm 0.01$ |
| BRCA1 (Bard1 binding) | $0.33 \pm 0.02$ | $0.32 \pm 0.01$ | $0.32 \pm 0.03$ | $0.33 \pm 0.03$ | $0.35 \pm 0.03$ |
| BRCA1 (E3 ligase activity) | $0.16 \pm 0.01$ | $0.23 \pm 0.03$ | $0.28 \pm 0.07$ | $0.33 \pm 0.05$ | $0.37 \pm 0.04$ |
| Beta Lactamase | $0.39 \pm 0.03$ | $0.69 \pm 0.01$ | $0.84 \pm 0.01$ | $0.88 \pm 0.01$ | $0.89 \pm 0.01$ |
| E4B (U-box domain) | $0.15 \pm 0.03$ | $0.23 \pm 0.03$ | $0.35 \pm 0.01$ | $0.46 \pm 0.03$ | $0.53 \pm 0.04$ |
| Hsp90 | $0.29 \pm 0.02$ | $0.54 \pm 0.01$ | $0.67 \pm 0.01$ | $0.70 \pm 0.02$ | $0.72 \pm 0.05$ |
| PSD95 (pdz3 domain) | $0.20 \pm 0.02$ | $0.46 \pm 0.05$ | $0.68 \pm 0.01$ | $0.76 \pm 0.01$ | $0.81 \pm 0.02$ |
| Pab1 (RRM Domain) | $0.42 \pm 0.07$ | $0.60 \pm 0.02$ | $0.76 \pm 0.01$ | $0.80 \pm 0.01$ | $0.83 \pm 0.02$ |
| Protein G | $0.34 \pm 0.15$ | $0.62 \pm 0.03$ | $0.85 \pm 0.02$ | $0.89 \pm 0.02$ | $0.93 \pm 0.01$ |
| Ubiquitin (E1 activation) | $0.41 \pm 0.06$ | $0.58 \pm 0.02$ | $0.67 \pm 0.02$ | $0.72 \pm 0.02$ | $0.74 \pm 0.02$ |
| Yap65 (WW domain) | | $0.38 \pm 0.06$ | $0.58 \pm 0.06$ | $0.68 \pm 0.05$ | $0.78 \pm 0.05$ |

**Table S5.** Aggregate spearman *ρ* measure across models and datasets. 34-layer Transformer pre-trained on UR50/S. For intraprotein models, the train/valid data was randomly partitioned five times. The mean ± standard deviation across the five runs is reported. No standard deviations are reported for LOPO experiments, as the evaluation is performed across all proteins.

| dataset<br>model | spearmanr | Envision (LOPO) | DeepSequence |
| --- | --- | --- | --- |
|  | Envision |  |  |
| Transformer | $0.71 \pm 0.20$ | 0.51 | $0.70 \pm 0.15$ |
| LSTM biLM (Large) | $0.65 \pm 0.22$ | | $0.62 \pm 0.19$ |
| Gray, et al. 2018 | $0.64 \pm 0.21$ | 0.45 | |
| Riesselman, et al. 2018 | | | $0.48 \pm 0.26$ |

**Figure S6.**
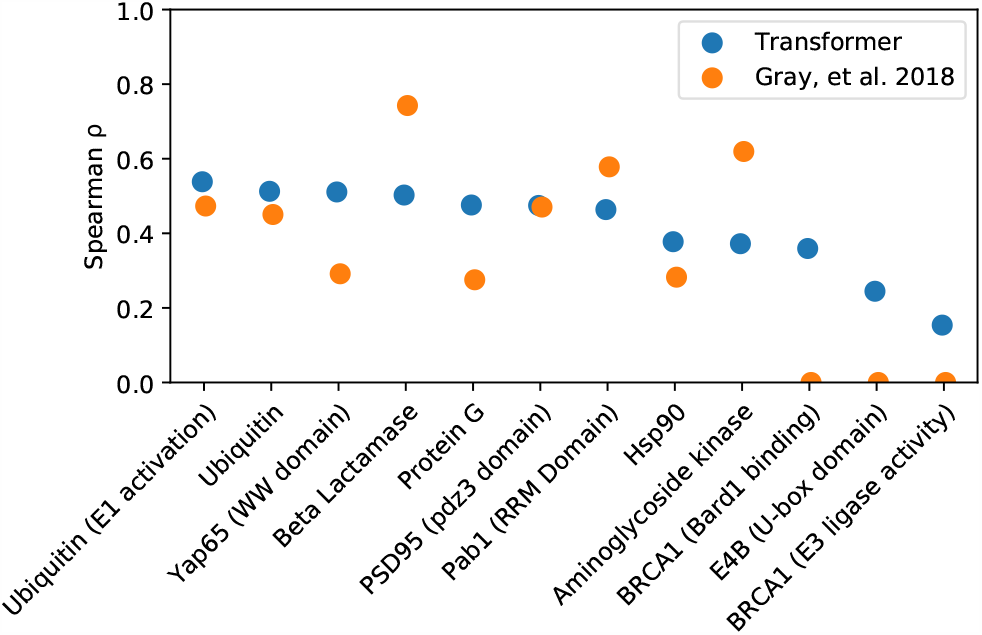
Leave-one-out experiment on Envision dataset (Gray et al., 2018). Pre-training improves the ability of the Transformer to generalize to the mutational fitness landscape of held-out proteins. All mutagenesis data from the protein selected for evaluation are held out, and the model is supervised with data from the remaining proteins. For each evaluation protein, a comparison is shown for the 34-layer Transformer pre-trained on UR50/S.

**Table S6.**
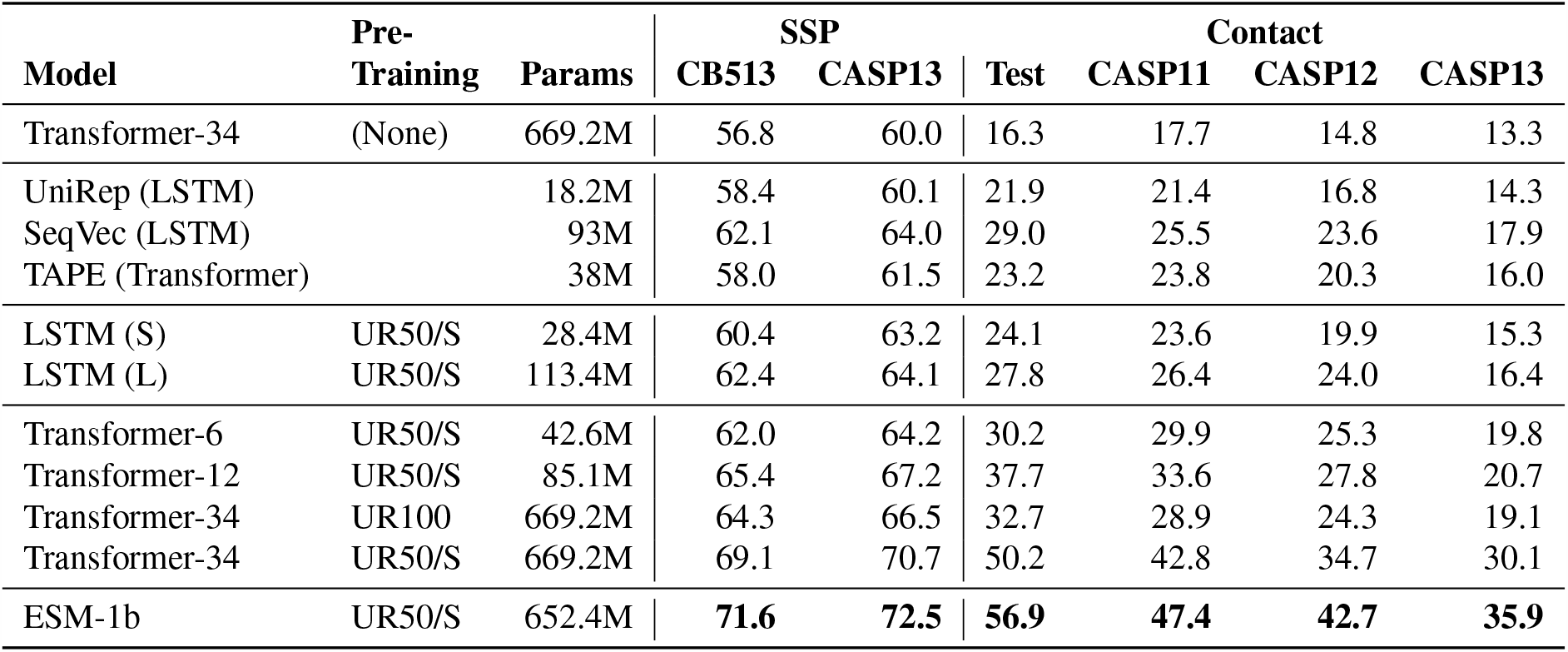
Comparison to related methods. Top-L long-range contact precision. Predictions are directly from protein sequence, no coevolutionary features or MSAs used. Test is RaptorX test set of Wang et al. (2017). Model weights for related work are obtained and evaluated in our codebase with same downstream architecture, training, and test data.

**Figure S7.**
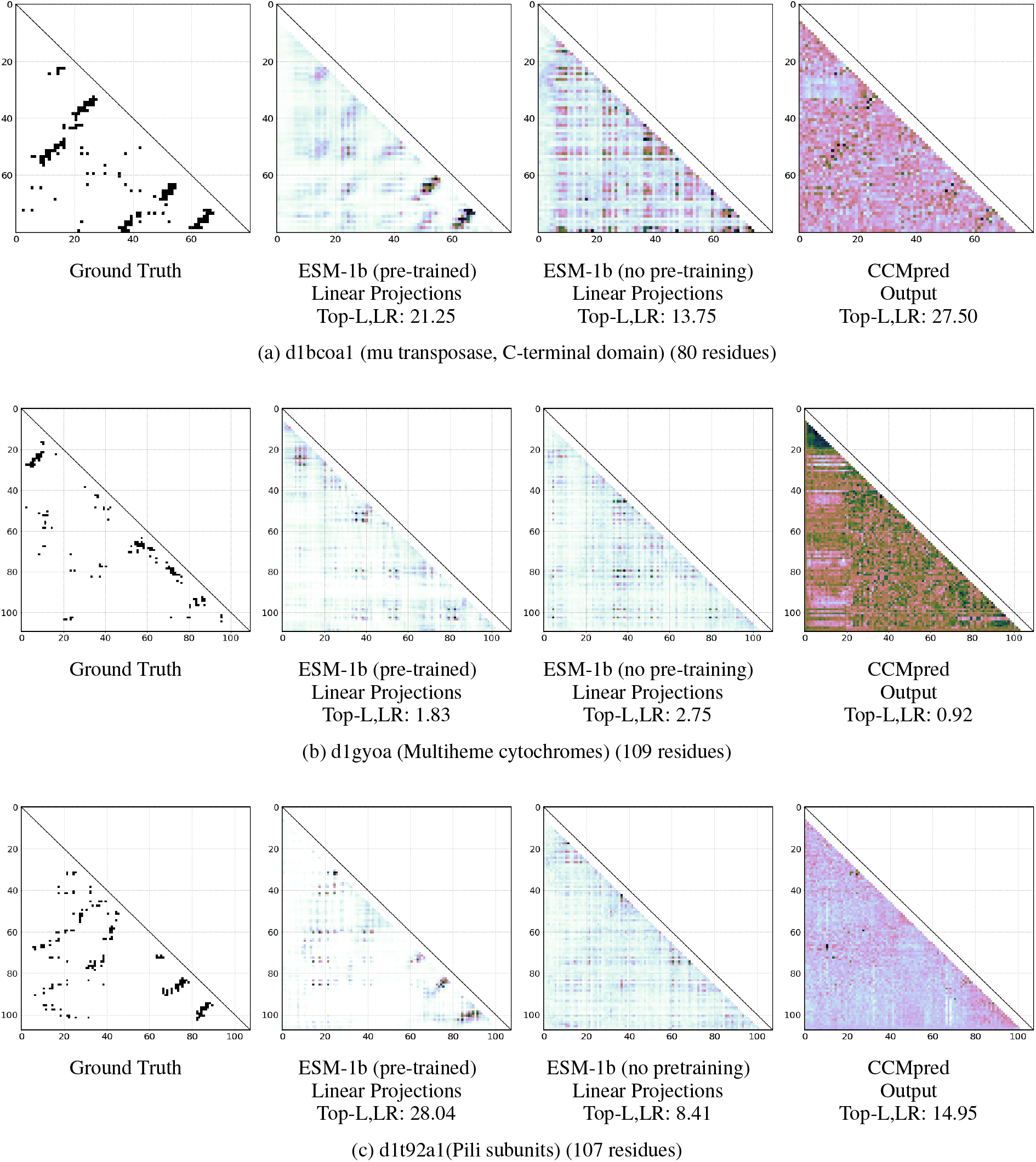
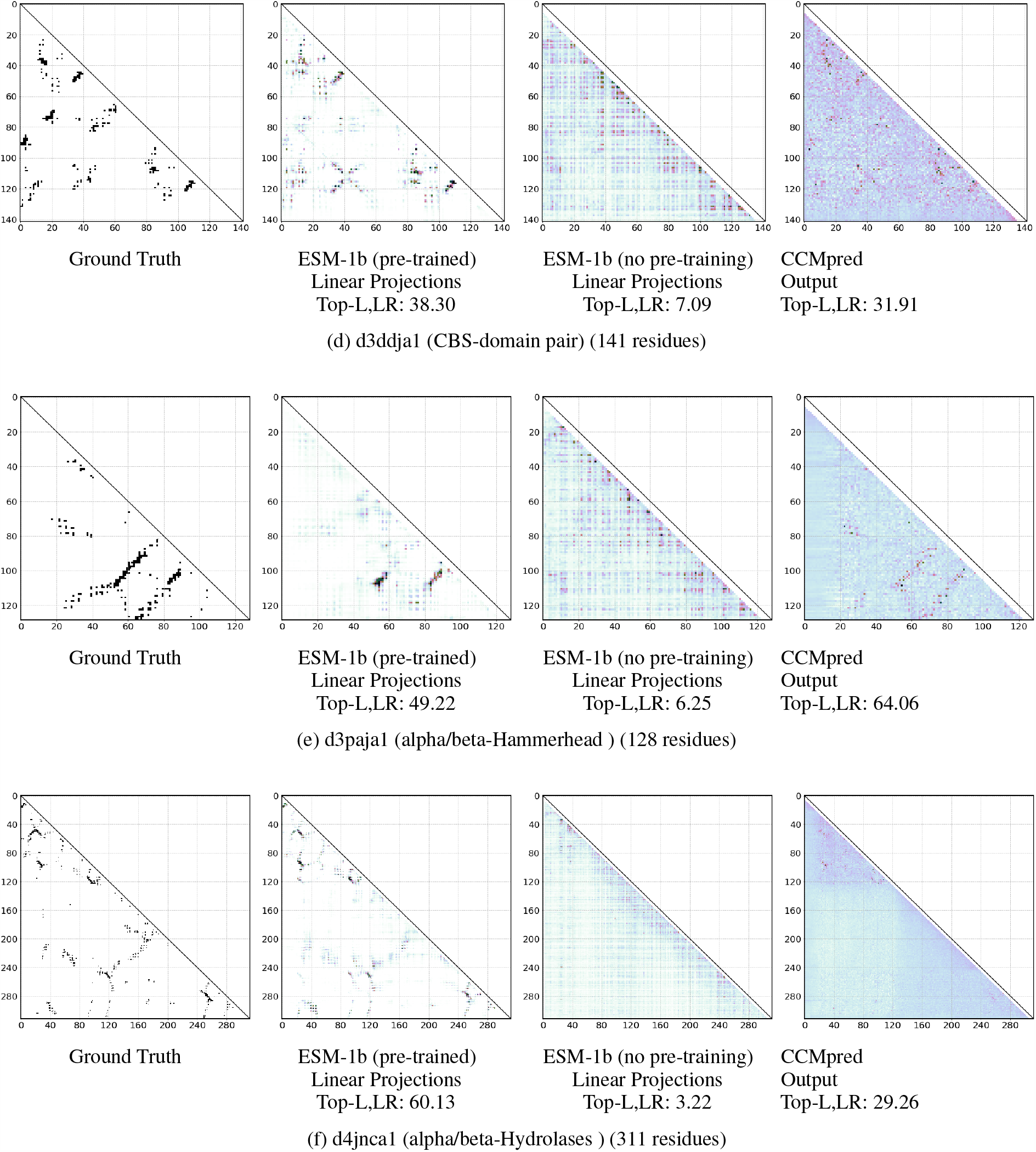
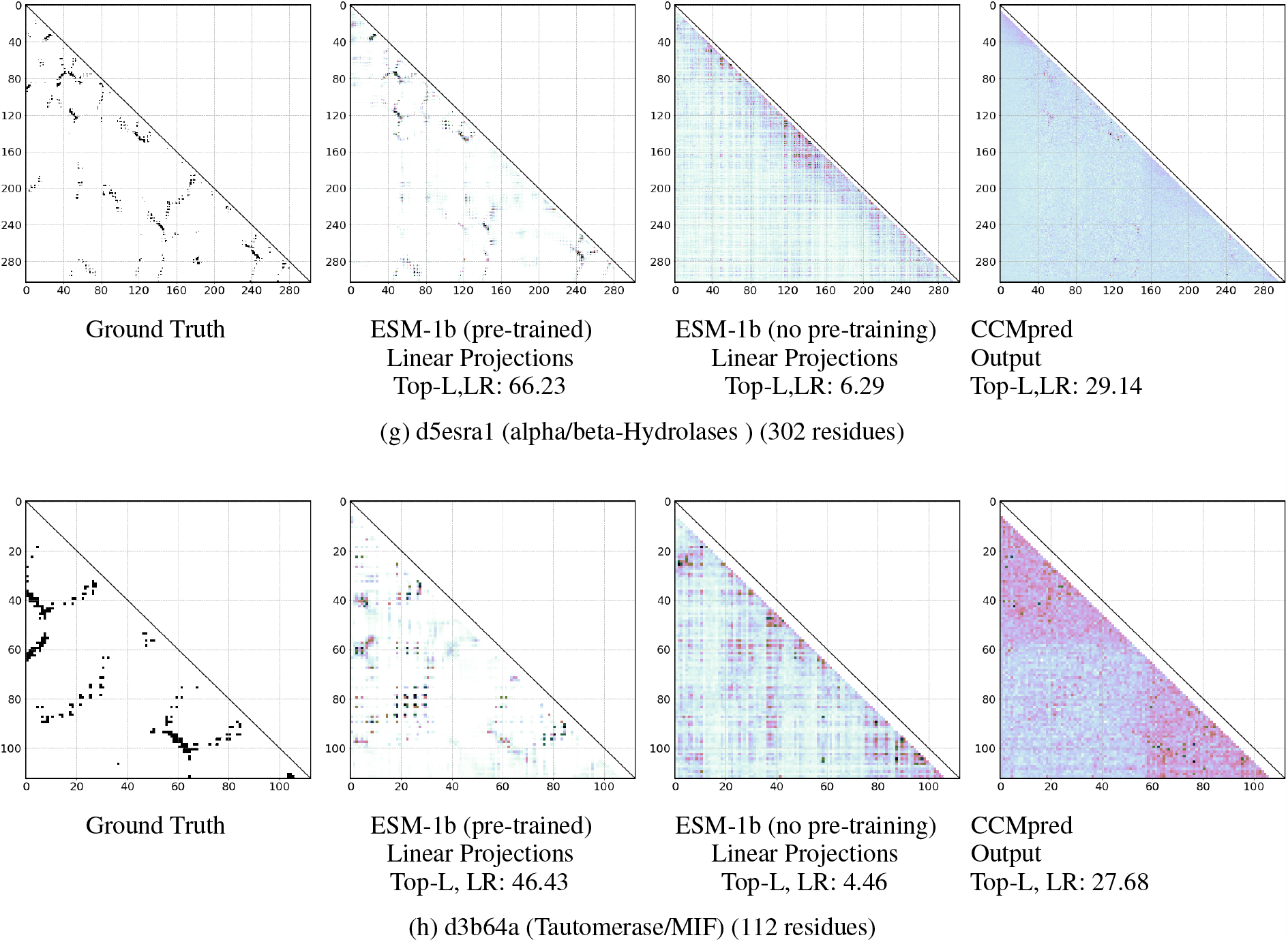
Comparison of ground truth contact map, projections from ESM-1b with and without pre-training, and CCMpred output. Labels indicate SCOPe domain, fold name, and number of residues. Eight domains randomly sampled from fold-level test sets are shown.

